# Comparative CRISPRi screens reveal a human stem cell dependence on mRNA translation-coupled quality control

**DOI:** 10.1101/2025.01.07.631643

**Authors:** Geraldine Rodschinka, Sergio Forcelloni, Andrew Behrens, Henrick Riemenschneider, Dieter Edbauer, Sascha Wani, Danny Nedialkova

## Abstract

The translation of mRNA into proteins in multicellular organisms needs to be carefully tuned to changing proteome demands in development and differentiation, and defects in translation often have a disproportionate impact in distinct cell types. Here we used inducible CRISPR interference screens to compare the essentiality of genes with functions in mRNA translation in human induced pluripotent stem cells (hiPSC) and hiPSC-derived neural and cardiac cells. We find that core components of the mRNA translation machinery are broadly essential, but the consequences of perturbing translation-coupled quality control factors are highly cell type-dependent. Human stem cells critically depend on pathways that detect and rescue slow or stalled ribosomes, and on the E3 ligase ZNF598 to resolve a novel type of ribosome collisions at translation start sites on endogenous mRNAs with highly efficient initiation. Our findings underscore the importance of cell identity for deciphering the molecular mechanisms of translational control in metazoans.

## Introduction

The human genome contains ∼20,000 predicted protein-coding genes, but only half are expressed at a time^1^, and fewer than a fifth are expressed at similar levels in all cell types^2^. In metazoans, intricately orchestrated developmental programs generate cell types with highly specialized functions and distinct protein content. This poses unique challenges to the mRNA translation machinery, which needs to accommodate rapid shifts in global and specific protein demands during developmental transitions^3^ and to faithfully synthesize even extremely long proteins such as titin (∼34,000 amino acids) in millions of copies in specialized cell types^4^. The initiation, elongation, and termination of protein synthesis by the ribosome is orchestrated by a multitude of factors that determine the accuracy and efficiency of each of these steps. Their tasks are complicated by variability in the untranslated regions (UTRs) that flank the same coding sequence (CDS) in mRNAs from different tissues, which can modulate protein output^5–7^. As the length of human proteins varies over 400-fold and their intracellular abundance spans six orders of magnitude^8^, quality control pathways mitigate errors in mRNA translation and dispose of problematic mRNAs and incomplete nascent chains that could be toxic to cells^9,10^. Failure of these surveillance mechanisms has been linked to human neurological disorders^11^.

The complexity of the mRNA translation machinery makes it challenging to determine how its plasticity is achieved during development, a task that is further compounded by the early embryonic lethality upon constitutive knockout of its components in mammals. Studies using conditional knockouts have shown that perturbations in mRNA translation can have outsized effects in distinct cell types and developmental stages^12–14^ but the underlying mechanisms have been a matter of debate^15,16^. The physiological signals that activate translation-coupled quality control pathways remain equally enigmatic, and their mechanistic dissection has so far relied on reporters mimicking problematic mRNAs^9,10^.

The function of human genes has been probed in high throughput in a range of immortalized and cancer-derived cell lines^17^, but these models are of limited value for dissecting cell type-specific regulation due to their genetic heterogeneity, unstable genomes, and aberrant gene expression^18,19^. Translational control, in particular, is extensively rewired to sustain the rapid proliferation of cancer cells^20^. Human induced pluripotent stem cells (hiPSC), by contrast, can self-renew without transformation and can give rise to nearly any cell type in culture^21^. Recent advances enable the robust differentiation of hiPSC along distinct cell lineages, as well as CRISPR-Cas9-based functional genomics in these models^22–24^. Here we took advantage of these technologies to define the essentiality of 262 core and regulatory genes from the mRNA translation machinery with CRISPR interference (CRISPRi) screens in hiPSC and a range of hiPSC-derived cell types. In contrast to the nearly universal essentiality of core ribosomal proteins and translation factors, the loss of many proteins that mediate translation-coupled quality control was detrimental only in some cell contexts while, surprisingly, being neutral or even advantageous in others. Such divergent genetic dependencies were especially pronounced in pathways that detect and rescue slow or stalled translating ribosomes, which we find are essential for growth and survival in human stem cells, but dispensable in the widely used immortalized HEK293 cell line. We identified distinct stress responses upon perturbing ribosome rescue pathways in different cellular contexts, and discovered a novel role for the E3 ligase ZNF598 in detecting ribosome collisions during translation initiation on endogenous mRNAs in human stem cells. Our study establishes an experimental platform for dissecting the cell context-dependent molecular determinants of fundamental biological processes.

## Results

### Comparative genetic screening in hiPSC-derived neural and cardiac cell types by inducible CRISPRi

To explore the rewiring of translational control as a function of cell identity, we profiled the reliance of different human cell types on components of the core and regulatory translational machinery by inducible CRISPRi screening^25,26^. This gene silencing approach combines a single guide RNA (sgRNA) targeting a promoter with catalytically inactive Cas9 (dCas9) fused to a transcriptional repressor domain (e.g. KRAB), which thus inhibits transcription^25^. Additionally, because it does not introduce double stranded DNA breaks, CRISPRi does not trigger the p53-mediated toxicity in human pluripotent stem cells, which is an obstacle to genetic screening^27^. We used a previously validated workflow^26^ to enable temporally controlled gene knockdown. Briefly, we inserted a doxycycline-inducible KRAB-dCas9 expression cassette at the *AAVS1* safe harbour locus in the reference hiPSC line *kucg-2*^28,29^ and in HEK293 for comparison (hereafter referred to as hiPSCi and HEK293i). The expression of the doxycycline-controlled reverse transcriptional activator (rtTA) is driven by the constitutive CAG promoter, whereas the expression of KRAB-dCas9 and mCherry (separated by a P2A ribosomal skipping sequence) are controlled by the doxycycline-response element TRE3G from the opposite DNA strand (**Fig. 1a**). This design prevents leaky expression in the absence of doxycycline^26^, and we confirmed that KRAB-dCas9 was undetectable in hiPSCi in the absence of doxycycline by western blotting and mass spectrometry (**Extended Data Fig. 1a**).

**Fig. 1:**
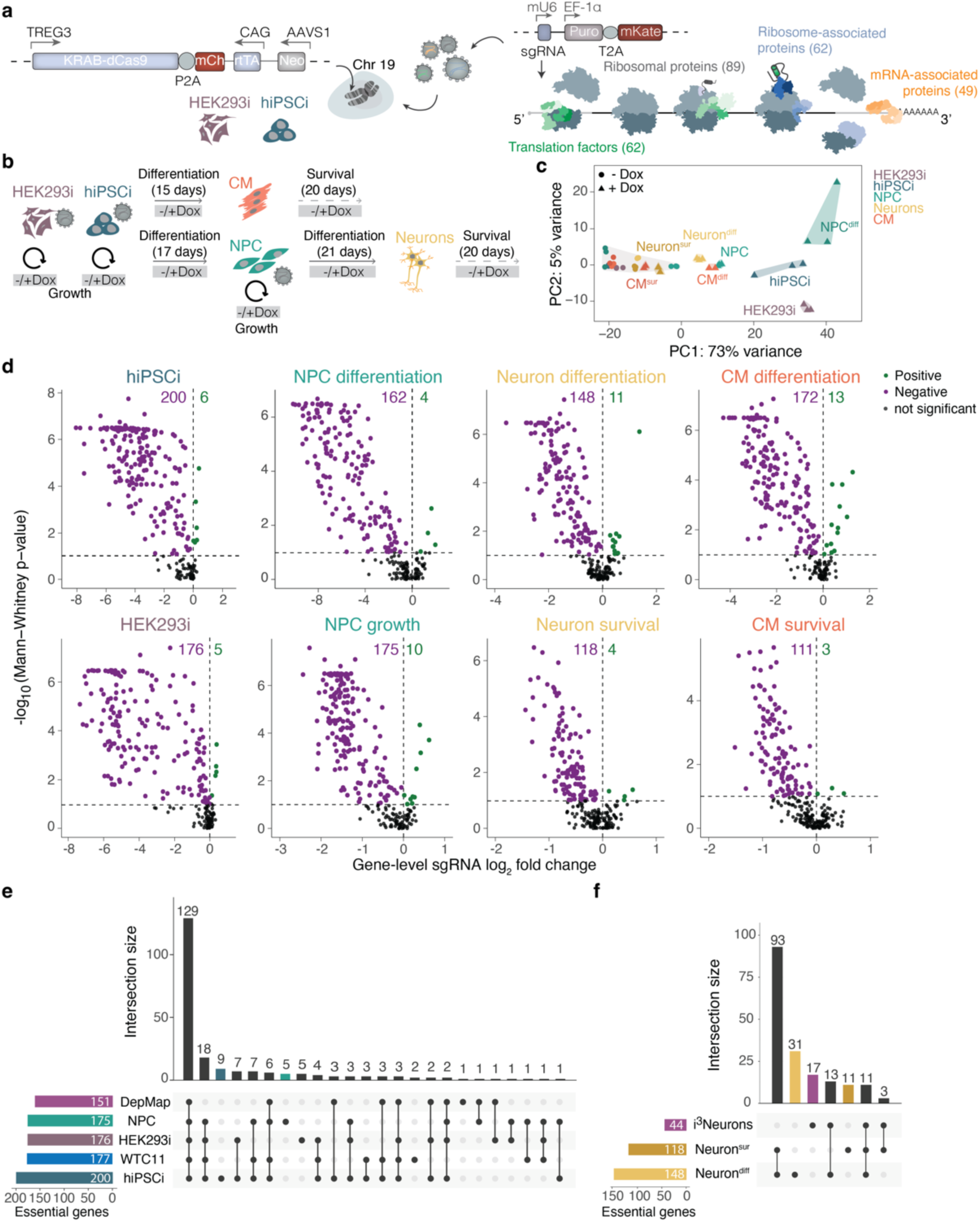
Comparative inducible CRISPRi screens identify essential components of the mRNA translation machinery in human cells. **a,** Schematic of inducible CRISPRi cell line generation and screening. Gene counts for major functional groups of the human translation machinery included in the sgRNA library are shown on the right. Neo: neomycin resistance gene; mCh: mCherry; Puro: puromycin resistance gene. **b,** Schematic of the workflow of hiPSCi differentiation into neural progenitor cells (NPC), neurons, and cardiomyocytes (CM) and the different types and duration of inducible CRISPRi screens. **c,** Principal component (PC) analysis of variance stabilizing-transformed count data for sgRNAs from DESeq2 (*n*=2 biological replicates for CM; *n*=3 biological replicates for hiPSCi, HEK293i, NPC, and neuron screens). Percent variance explained by each PC shown in parentheses; diff, differentiation; sur, survival. **d,** Volcano plots of gene-level sgRNA log_2_ fold change (FC) (mean of the top 3 sgRNA per gene by magnitude) and Mann-Whitney *p-*values (from comparisons of all sgRNAs targeting a given gene to all negative control sgRNAs) for each screening condition relative to matched uninduced (-Dox) controls. Genes with significant (Mann-Whitney test *p* ≤ 0.1) positive or negative enrichment scores are shown in green and purple, respectively. **e,f,** UpSet plots showing overlap of genes with significant (Mann-Whitney test *p* ≤ 0.1) negative enrichment scores in (**e**) hiPSCi, HEK293i, and NPC screens in comparison to common essential genes in cancer cell lines (DepMap^17^) and genome-wide CRISPRi in the WTC11 hiPSC line^34^, and (**f**) neuron differentiation (Neuron^diff^) or survival (Neuron^sur^) screens compared to a genome-wide CRISPRi screens in WTC11-derived i^3^Neurons^35^.

We designed a pool of sgRNAs targeting the promoters of 262 human genes encoding ribosomal proteins, core translation factors, cytoplasmic mRNA stability factors, proteins interacting with nascent chains, mRNA and ribosome quality control factors, and 9 cell-specific marker genes as controls (**Fig. 1a**). We used CRISPRiaDesign^30^ to design a focused sgRNA library containing 9 sgRNAs per annotated TSS in our target gene set, retaining sgRNAs with the highest predicted activity scores and no predicted off-targets. The resulting library (3000 sequences, 10% non-targeting controls) was cloned in a mouse U6 promoter-based lentiviral expression vector (**Fig. 1a**).

We used established protocols for the robust differentiation of hiPSCi into neural progenitor cells (NPC), neurons, and cardiomyocytes (CM)^29^ (**Fig. 1b**), which comprise cell types and developmental transitions subject to extensive translational control^3,31–33^. Lineage-specific markers were uniformly expressed in hiPSCi (*NANOG*, *POU5F1*) and their differentiated counterparts (*PAX6* and *NES* in NPC, *CHAT* and *MAP2* in neurons, and *CTNT* and *ACTN2* in CM)^29^ (**Extended Data Fig. 1b**), confirming the purity of these cultures. Analysis of mCherry levels by flow cytometry as a measure of doxycycline response revealed a robust expression of KRAB-dCas9 after induction in HEK293i, hiPSCi, NPC, neurons, and CM (**Extended Data Fig. 1c)**.

To compare gene essentiality across cellular contexts and cell state transitions, hiPSCi, NPC, and HEK293i were transduced with the lentiviral sgRNA library while ensuring that only one sgRNA is expressed in each cell. Following puromycin selection to eliminate untransduced cells, matched cultures grown without or with doxycycline were collected after 10 population doubling times (**Extended Data Fig. 1d**) to conduct growth screens. We also conducted differentiation and survival screens by assessing gene essentiality during or after the derivation of NPC and CM from transduced hiPSC, and neurons from transduced NPC (**Fig. 1b**). Principal component analysis of sgRNA counts revealed a clustering of all uninduced controls, as well as biological replicates of individual screens (**Fig. 1c**). The most pronounced changes in sgRNA frequency were in growth screens for hiPSCi and HEK293i and surprisingly also during NPC differentiation. These data indicate that the transition of hiPSC to NPC is particularly sensitive to perturbations in mRNA translation.

We next calculated gene-level enrichment or depletion scores with an established CRISPRi screen analysis pipeline^30^, which averages the log_2_ fold change (FC) of the top three sgRNAs targeting each gene TSS by magnitude and calculates a Mann-Whitney *p*-value using all sgRNAs targeting the same gene compared to non-targeting control sgRNAs. This analysis revealed between 111 and 200 targets as significantly depleted (*p* ≤ 0.1) and 3 to 13 targets as significantly enriched in the different screens (**Fig. 1d**). sgRNAs targeting cell markers were depleted only in the relevant screens (**Extended Data Fig. 1e**), supporting our ability to detect cell context-specific gene essentiality. Among our 262 targets, we identified 150 out of the 151 genes (99%) that scored as “common essential” in hundreds of human cancer cell lines (DepMap)^17^, and 175 out of 177 (99%) negative hits from a genome-wide CRISPRi screen in the WTC11 hiPSC line^34^ (**Fig. 1e**). 27 genes (15%) were essential in our *kucg-2*-derived hiPSCi but not in WTC11, which may reflect differences in genetic background, screen setup, or a higher sensitivity of focused versus genome-wide screens. We identified 148 genes essential for neuronal differentiation and 118 genes essential for neuron survival (**Fig. 1f**), while only 44 of our 262 targets had previously scored as essential in a genome-wide screen in WTC11-derived i^3^Neurons^35^, 24 of which (55%) we recovered as well. Our inducible CRISPRi screens thus recapitulate known genetic dependencies and identify novel ones in hiPSC and their derived cell types.

A comparison across all screening contexts revealed that hiPSCi had the highest sensitivity to mRNA translation perturbations, with 200 out of 262 (76%) genes scoring as essential compared to 176 and 175 (67%) in HEK293i and NPC, respectively (**Extended Data Fig. 1f**). This could be in part be linked to the exceptionally high global protein synthesis rates in hiPSCi (**Extended Data Fig. 1 g,h**). The cell state transitions we queried (hiPSCi→NPC, hiPSCi→CM, and NPC→neurons) were also highly vulnerable to the loss of mRNA translation factors (162, 172, and 148 essential genes, respectively). The lower overall number of hits in the neuron and CM survival screens may, at least in part, be due to less efficient protein depletion in non-dividing cells because of lack of dilution by cell division^36^. Notably, genetic dependencies specific for a single cell type were extremely rare: only one gene essential for the survival of neurons (*NAA11*) or CM survival (*CPEB2*) and four genes essential for HEK293i growth (*CARHSP1*, *EIF4E3*, *EIF4G3*, and *IGF2BP2*) did not score as hits in any other of our screening contexts.

To validate our screen results, we selected the two most highly effective sgRNAs (sgRNA1 and sgRNA2) targeting 16 genes with differential essentiality (**Extended Data Fig. 2a**), and transduced them individually in hiPSCi, NPC, and HEK293i (**Extended Data Fig. 2b**). sgRNA enrichment or depletion in these individual experiments were highly correlated with the scores for the same sgRNAs in the screens (Spearman’s R=0.85 for hiPSCi, 0.72 for NPC and 0.51 for HEK293i; **Extended Data Fig. 2c**), and quantitative RT-PCR confirmed efficient knockdown for all targets in all three cell lines (**Extended Data Fig. 2d**). The differential importance of the proteins encoded by these 16 genes was not linked to their abundance in specific cell contexts, as most were present at similar levels in hiPSCi, NPC, and HEK393i based on quantitative mass spectrometry (**Extended Data Fig. 2e**). As human stem cells have higher proteasome activity than somatic cells^37^, we considered that other cell types may appear more robust to gene repression due to less efficient protein depletion. Immunoblot analysis of four target gene products revealed that they all decreased substantially in hiPSCi, NPC, and HEK293i (**Extended Data Fig. 2f**) and their depletion efficiency did not correlate with their essentiality in these cell contexts (**Extended Data Fig. 2a**). Collectively, these data suggest that the differential robustness to genetic perturbations we identified among cell types reflects unique demands on their protein synthesis machinery.

### Broad essentiality of ribosomal proteins and core translation factors

To determine how perturbations in distinct functional modules of the mRNA translation machinery impact cell growth, differentiation, and survival, we broadly grouped gene targets into ribosomal proteins (r-proteins), translation factors (initiation/elongation/termination), mRNA-associated proteins, and ribosome-associated proteins. A comparison of gene-level sgRNA log_2_ FC among subgroups revealed strong negative effects of targeting nearly all canonical r-proteins and translation factors in dividing cells (hiPSCi, NPC, HEK293i) and during differentiation (**Fig. 2a**). 76 out of the 79 canonical r-protein genes were essential in dividing cells, and 63 were essential across all cell contexts (**Fig. 2 a,b**). These data were consistent with the core essentiality of these genes in DepMap^17^ and in genome-wide CRISPRi screens in WTC11 hiPSC and the H1 human embryonic stem cell (hESC) line^34^. Depletion of factors involved in ribosome subunit export and cytosolic ribosome maturation (*LSG1*, *LTV1* and *RIOK2*), which we grouped with ribosome-associated proteins also had concordant strongly negative effects in all screens (**Fig. 2b**). Together, these data indicate that new ribosomes must be continuously produced to maintain translational homeostasis also in non-dividing cell types.

**Fig. 2:**
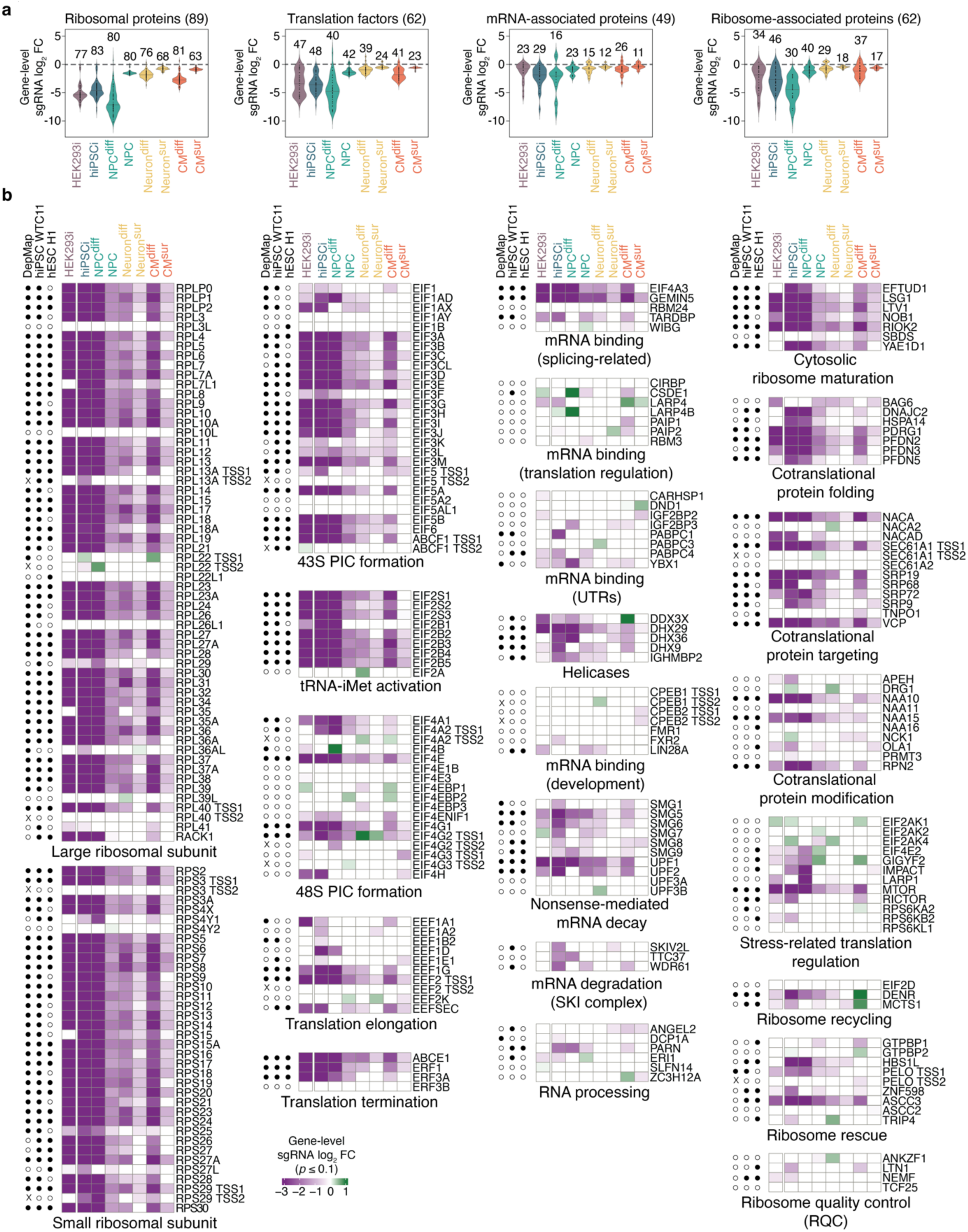
Common and cell type-specific effects of mRNA translation perturbations in human cells. **a,** Violin plots of gene-level sgRNA log_2_ fold changes (mean of the top 3 sgRNA per gene by magnitude) in each screen separated into functional gene groups (ribosomal proteins, translation factors, mRNA-associated proteins, ribosome-associated proteins). The number of genes with significant (Mann-Whitney *p* ≤ 0.1) enrichment or depletion in each screen are indicated above each violin. **b,** Heatmaps of data in **(a**). Significant (Mann-Whitney *p* ≤ 0.1) gene-level enrichment or depletion in each screen indicated by colour; white: not significant in the respective screen. Data for genes with more than one annotated transcription start site (TSS) were analysed separately. Open and closed circles indicate non-essential and essential genes, respectively, in the common essential gene set from DepMap 23Q4^17^ and genome-wide CRISPRi screens in WTC11 hiPSC and H1 hESC^34^; x indicates the absence of data for alternative TSSs in DepMap.

Two of the small ribosomal subunit (40S) and seven of the large ribosomal subunit (60S) r-protein genes have paralogs, some of which have been proposed to endow ribosomes with distinct functions in specific tissues^38,39^. Depletion of *RPL22* or its paralog *RPL22L1* had minimal effects in all screens (**Fig. 2b**), consistent with their functional redundancy and the compensatory expression of *Rpl22l1* in *Rpl22^−/−^* mice^40^. Among the remaining non-canonical r-protein paralogs, *RPS27L* and *RPL36AL* scored as essential in a subset of our screens (**Fig. 2b**), suggesting that their protein products function in selected cell types.

Our screens also revealed divergent effects upon repression of gene paralogs for canonical initiation and elongation factors. Knockdown of *EEF1A2,* the brain- and muscle-specific paralog of the ubiquitously expressed elongation factor-encoding *EEF1A1* gene^41^, inhibited neuron survival (**Fig. 2b**). Repression of the initiation factor eIF4G1 paralog *EIF4G2* inhibited cell proliferation but surprisingly promoted neuron differentiation and survival (**Fig. 2b**). The depletion of proteins that globally repress translation initiation downstream of growth or stress signalling (eIF4EBPs or EIF2AK) similarly promoted cell proliferation or survival in specific contexts (**Fig. 2b**). Collectively, these findings demonstrate the ability of our comparative screening approach to identify common and cell context-specific genetic dependencies.

### Human stem cells are highly sensitive to perturbations in translation-coupled quality control

In contrast to the broad essentiality of proteins that mediate the core steps of mRNA translation, we found surprisingly divergent genetic dependencies for many translation-coupled quality control pathways. hiPSCi were highly sensitive while HEK293i were highly resilient to depletion of the SKI complex, which directs cotranslational mRNA degradation by the exosome^42,43^, and *DNAJC2* and *HSPA14*, whose protein products form the mammalian ribosome-associated complex that assists nascent chain folding (**Fig. 2b**)^44^. Knockdown of *GIGYF2* and *EIF4E2*, whose products form a repressor complex that limits translation initiation on faulty mRNAs^45,46^, inhibited hiPSCi proliferation, but enhanced NPC growth (**Fig. 2b**). Another heterodimeric complex encoded by *DENR* and *MCTS1* that regulates ribosome recycling^47,48^ was essential in dividing cells, but its loss surprisingly promoted CM differentiation (**Fig. 2b**).

Factors mediating the rescue of ribosomes that stall or collide when translating problematic mRNA or peptide sequences^9^ had the most divergent essentiality in our screens. PELO and its partner HBS1L destabilize ribosomes stalled at the 3’ end of truncated mRNA reporters^49–53^. Depletion of PELO or HBS1L strongly inhibited the growth of hiPSCi and neural differentiation. While HEK293i were also moderately sensitive to PELO loss, they were surprisingly robust to HBS1L depletion (**Fig. 2b**), indicating that the two proteins may function independently in some cellular contexts^13,54^. In line with this hypothesis, *PELO* scored as a common essential gene in cancer cell lines as well as in WTC11 hiPSC and H1 hESC, while *HBS1L* was only essential in WTC11 hiPSC and H1 hESC^34^ (**Fig. 2b**).

Unresolved ribosome stalling can lead to ribosome collisions, which trigger distinct quality control and stress response pathways^55^. Collided ribosomes are thought to be recognized by the E3 ligase ZNF598, which ubiquitinates the 40S r-proteins uS10 and eS10^56–61^ to enable ribosome disassembly by the helicase ASCC3^62,63^. While *ASCC3* was near-universally essential, *ZNF598* repression was highly detrimental for hiPSCi and to a lesser extent NPC growth, it was better tolerated by HEK293i (**Fig. 2b**), suggesting that ASCC3 may not always function downstream of ZNF598. Notably, *ZNF598* also scored as essential in genome-wide CRISPRi screens in WTC11 hiPSC and H1 hESC^34^ (**Fig. 2b**). Not all factors implicated in ribosome rescue were essential in human stem cells, however: depletion of N4BP2, the homolog of yeast Cue2 and worm NONU-1 that cleave mRNAs in the A site of collided ribosomes^64,65^, or EDF1, which binds collided ribosomes^66,67^, had no appreciable effect on either hiPSCi or HEK293i growth (**Extended Data Fig. 2g,h**). By contrast, perturbing the ribosome quality control (RQC) pathway that disposes of nascent chains on stalled ribosomes through NEMF recruitment of the E3 ligase LTN1^68–70^ was also selectively detrimental in hiPSCi, as well as in WTC11 hiPSC and H1 hESC^34^ (**Fig. 2b**). Collectively, these data indicate that the functional importance of translation-coupled quality control strongly depends on the cellular context, and is disproportionally high in human stem cells.

### Cellular robustness to perturbed ribosome rescue is not due to functional redundancy

We next asked whether the selective robustness of cells to defective ribosome rescue is linked to potential functional redundancies in specific contexts. To test this, we asked how the repression of *ZNF598*, *ASCC3*, *PELO*, or *HBS1L* influences ribosome stalling in different cell types. In individual sgRNA experiments, the knockdown of these four genes with one of the two most highly effective sgRNAs from the screens was highly detrimental for hiPSCi growth and survival, while it was much better tolerated by HEK293i cells (**Fig. 3 a,b; Extended Data Fig. 2i**). Since the *HBS1L* locus also encodes SKI7, a cytoplasmic exosome cofactor is expressed from a minor alternative splice isoform^71,72^, we asked if the essentiality of *HBS1L* in hiPSCi could be partially attributed to SKI7 depletion. However, re-introduction of cDNA encoding wild-type but not GTPase-deficient *HBS1L* rescued the growth defects resulting from *HBS1L* knockdown in hiPSCi (**Extended Data Fig. 2j**). These findings indicate that the GTPase activity of HBS1L is essential for human stem cell proliferation and survival.

**Fig. 3:**
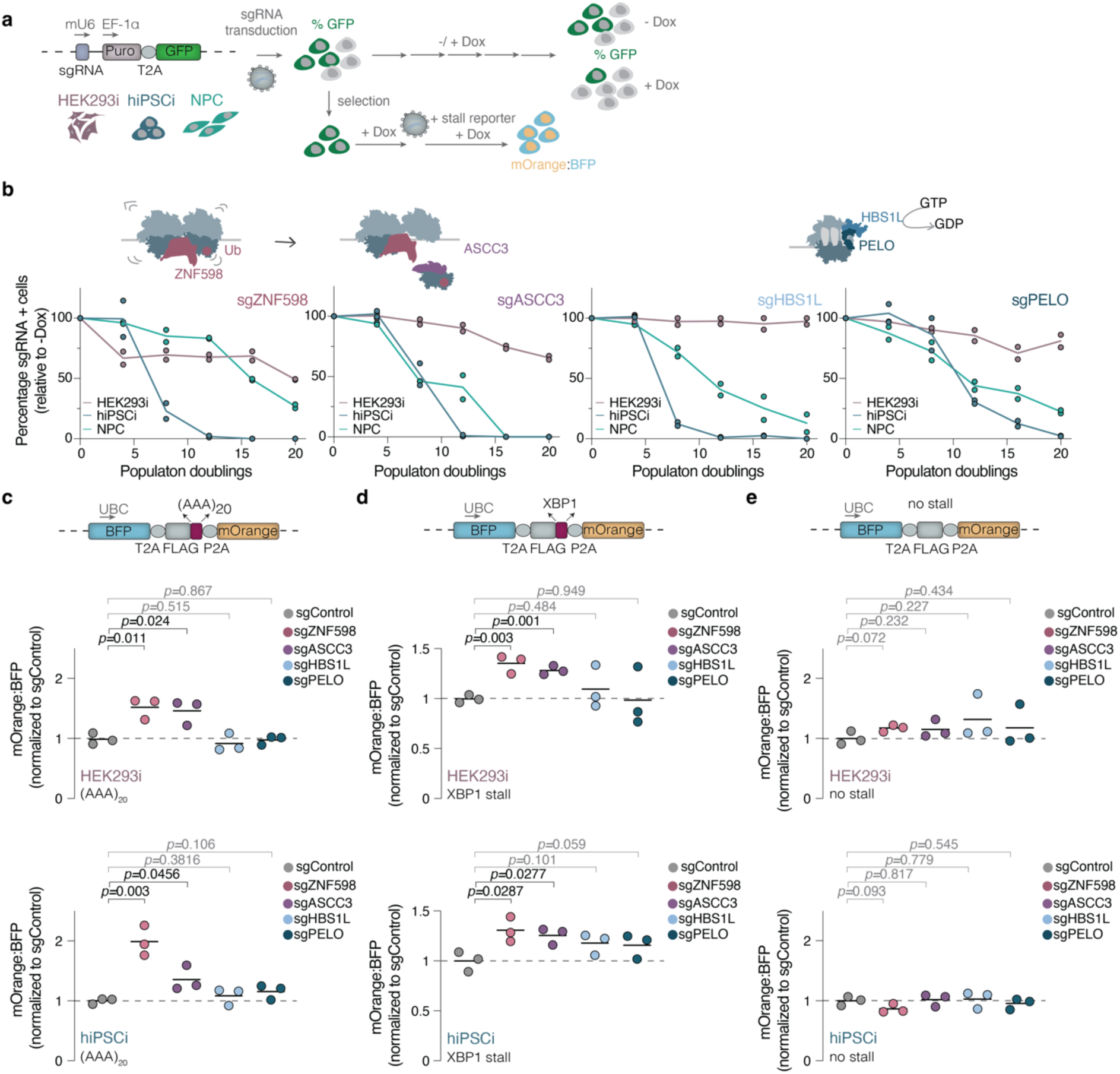
Cellular robustness to perturbed ribosome rescue is not due to functional redundancy. **a,** Schematic of expression constructs and workflows for growth competition and stalling reporter readthrough assays. **b,** Growth assays of cells transduced with the most potent sgRNA targeting *ZNF598*, *ASCC3*, *HBS1L*, or *PELO* in the hiPSCi screen (*n=*2 biological replicates; line: average). The percentage of GFP-positive (GFP+) cells was measured by flow cytometry (>10,000 cells/ analysis) every four population doublings and normalized to GFP+ cell numbers in matched uninduced (-Dox) controls. **c,d,e,** Stalling readthrough of reporters containing an AAA-encoded stretch of twenty lysines (AAA_20_) (**c**) the XBP1 arrest peptide (**d**) and a no-stall control (**e**). The median fluorescence intensity for BFP and mOrange was quantified by flow cytometry (>20,000 cells/ analysis) in the indicated cell lines transduced with the most potent sgRNA targeting *ZNF598*, *ASCC3*, *HBS1L*, or *PELO* based on the hiPSCi screen. The ratio of mOrange to BFP in knockdown cells was normalized to that for the same reporter in cells transduced with a non-targeting sgRNA (sgControl; *n*=3 biological replicates; *p*-values from unpaired two-tailed t-test).

Since the endogenous translation events targeted by ZNF598, ASCC3, PELO, or HBS1L in human cells are poorly defined, we measured the readthrough efficiency of model ribosome stalling substrates with an established fluorescence-based quantitative reporter assay^58,73^. The reporter consists of a BFP gene separated from an mOrange gene by a FLAG tag, which is flanked by 2A sequences that induce peptide bond formation skipping. Insertion of a ribosome stalling sequence downstream of FLAG results in a constant BFP production and variable mOrange levels dependent on stalling readthrough. We analysed the ratios of mOrange to BFP produced from reporters containing one of two well-characterized ribosome stalling sequences: an AAA-encoded stretch of twenty lysines (AAA)_20_, which mimics ribosome encounters with prematurely polyadenylated mRNA^58,73^, or the XBP1 arrest peptide, which tethers ribosomes translating the *XBP1-u* mRNA to the ER, poising it for intron excision upon ER stress to enable the production of the XBP1-s transcription factor^62,74–76^. In line with previous reports^63,73^, ZNF598 or ASCC3 depletion increased the readthrough of AAA_20_ and XBP1 stalls in both hiPSCi and HEK293i (**Fig. 3 c-e**), indicating that the higher robustness of HEK293i to the loss of these two factors is not due to functional redundancy.

### Ribosome rescue prevents a cytotoxic ISR in human stem cells

To define the mechanisms underlying the human stem cell-specific growth defects upon ribosome rescue perturbations, we next examined how these perturbations impact global translation rates and cell viability. As a positive control, we depleted the universally essential initiation factor EIF2S1 (**Extended Data Fig. 3a**), which reduced *de novo* protein synthesis in both HEK293i and hiPSCi by >80%. Strikingly, depletion of ZNF598, ASCC3, HBS1L, or PELO reduced *de novo* protein synthesis by 20-40% in hiPSCi, but had no significant effect in HEK293i (**Fig. 4a**). Lactate dehydrogenase (LDH) levels in culture medium, a proxy for membrane damage due to cytotoxicity^77^, were also significantly elevated upon knockdown of ribosome rescue factors in hiPSC but not in HEK293i (**Fig. 4b**). These data indicate that perturbed ribosome rescue induces cytotoxicity selectively in the stem cell context.

**Fig. 4:**
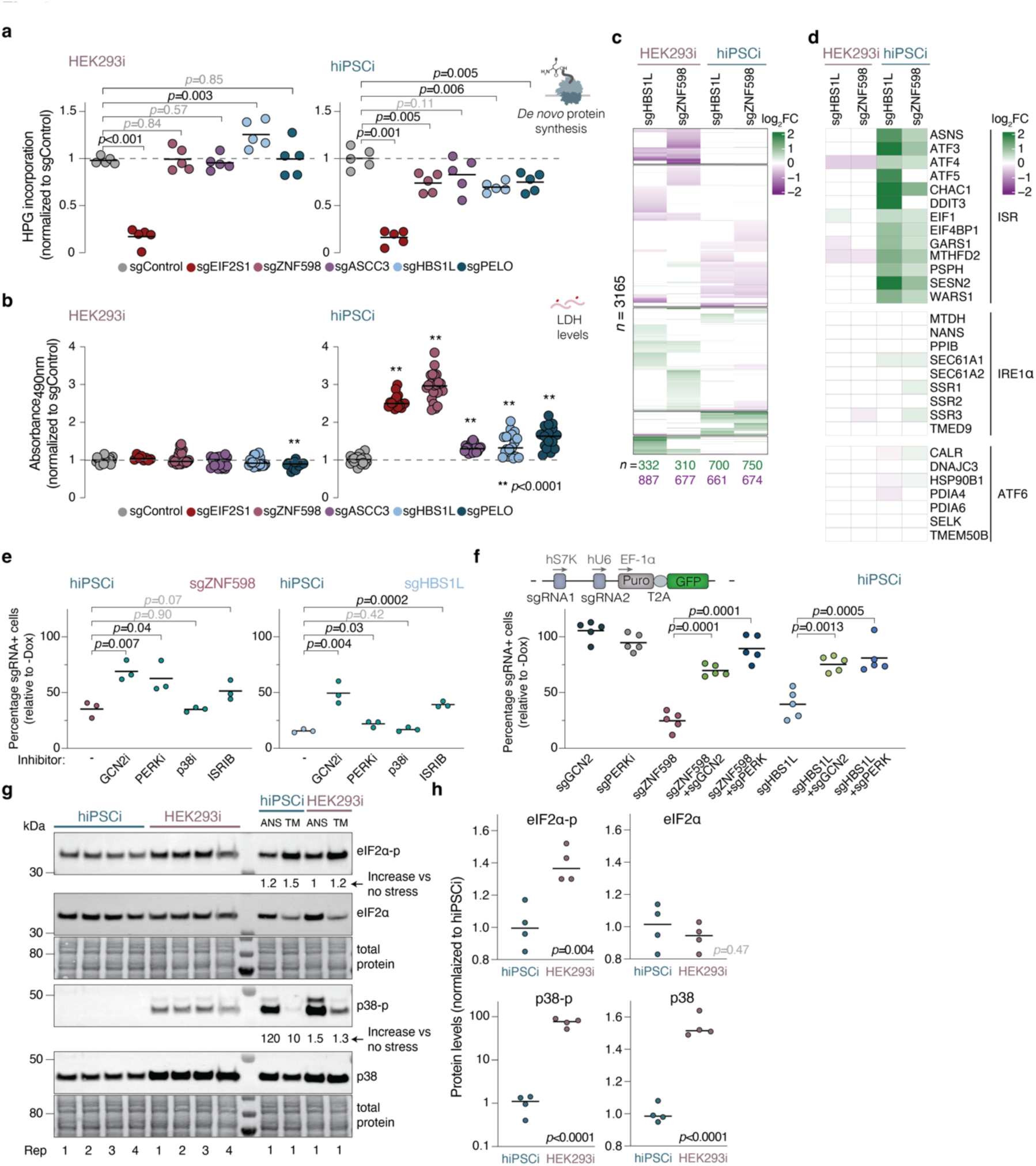
Ribosome rescue prevents a cytotoxic ISR in human stem cells. **a**, Global protein synthesis measurements in knockdown hiPSCi or HEK293i by L-Homopropargylglycine (HPG) labeling for 30 minutes. Median fluorescence intensity was quantified by flow cytometry (>10,000 cells/ analysis) and normalized to values from cells transduced with a non-targeting sgRNA (sgControl; *n=*5 biological replicates; *p*-values from unpaired two-tailed t-test). **b**, Lactate dehydrogenase (LDH) measurements in culture supernatants from knockdown hiPSCi or HEK293i. Values were normalized to supernatant from cells transduced with a non-targeting sgRNA (sgControl; *n=*4 technical replicates and ≥ 3 biological replicates; *p*-values from unpaired t-test). **c**, Heatmap of mRNAs differentially expressed upon *ZNF598* or *HBS1L* repression in hiPSCi or HEK293i (*n=*2 biological replicates; Benjamini–Hochberg-adjusted Wald test; *p-adj* ≤ 0.05; *n=*3165). **d**, Heatmap of a data subset from (**c**) showing genes upregulated downstream of ATF4 within the integrated stress response (ISR), or downstream of ATF6 or IRE1α upon ER protein folding perturbations^80^. **e,** Growth assays of hiPSCi expressing a sgRNA targeting *ZNF598* (day 6) or *HBS1L* (day 8) in the absence (-) or presence of inhibitors of GCN2 (GCN2i, A-92, 1.25 µM), PERK (PERKi, GSK2606414, 4 nM), p38 (p38i, SB203580, 1 µM), or the ISR (ISRIB, 50 nM) in comparison to uninduced controls (*n*=3 biological replicates, *p*-values from unpaired two-tailed t-test). **f,** Growth assays of hiPSCi expressing a sgRNA targeting *ZNF598*, *HBS1L*, *GCN2* and *PERK* alone or in combination in comparison to uninduced controls (*n*=5 biological replicates, *p*-values from unpaired two-tailed t-test). **g,** Immunoblot analysis of eIF2α phosphorylation, p38 phosphorylation, and total eIF2α and p38 levels in hiPSCi and HEK293i cells (*n*=4 biological replicates). hiPSCi treated with 2.5 µM tunicamycin (TM) for 2 hours served as a positive control for induction of eIF2α phosphorylation; hiPSCi treated with 0.05 mg/L anisomycin (ANS) for 15 min served as a positive control for induction of p38 phosphorylation. **h,** Quantification of signal intensity in **(g)** by densitometry (*p*-values from unpaired two-tailed t-test).

We next profiled the gene expression changes induced by knockdown of *HBS1L* or *ZNF598* by RNA-Seq. The patterns of gene expression changes were strikingly similar among the two genetic perturbations, but differed substantially between the two cell lines (**Fig. 4c**). hiPSCi experienced upregulation of genes with Gene Ontology (GO) terms related to chromosome organization, RNA processing and transport, and protein folding and localization (**Extended Data Fig. 3b**), whereas HEK293i exhibited downregulation of many genes with the same GO terms (**Extended Data Fig. 3c**). In hiPSCi but not in HEK293i, knockdown of either *ZNF598* or *HBS1L* resulted in a marked upregulation of integrated stress response (ISR) marker genes (**Fig. 4d**).

In mammals, the ISR is activated by one of four kinases: GCN2, PERK, PKR, or HRI, which sense a broad and partially overlapping range of stresses and phosphorylate the α subunit of eIF2^78^. This leads to a global repression of cap-dependent mRNA translation but it selectively increases *ATF4* translation by relieving the inhibitory effect of upstream open reading frames (uORFs) in its 5’ UTR. ATF4 is a transcription factor that induces a specific gene expression program to restore cellular homeostasis or trigger cell death^78,79^. ATF4 target genes^79,80^ were significantly upregulated upon depletion of ZNF598 and HBS1L in hiPSCi, while the expression of other stress response genes downstream of other stress sensors e.g. for protein misfolding in the ER (ATF6 and IRE1α^80^) was largely unchanged (**Fig. 4d**). In line with the ATF4-dependent gene expression signatures and the global reduction of protein synthesis (**Fig. 4a**), knockdown of *ZNF598* or *HBS1L* increased eIF2α phosphorylation levels in hiPSCi measured by flow cytometry (**Extended Data Fig. 3d**). Consistent with the lack of global translation defects or ISR gene expression signatures in HEK293i depleted for ZNF598 (**Fig. 4a, d**), no increase in eIF2α phosphorylation levels was observed upon knockout of *ZNF598* in HEK293T cells in previous work^57^.

Apart from the ISR, unresolved ribosome stalling and collisions can trigger the ribotoxic stress response (RSR) through phosphorylation of the MAP kinase p38 by ZAKα^55,81^. The levels of phosphorylated p38 were increased in hiPSCi depleted of ZNF598 or HBS1L, albeit to a variable extent (**Extended Data Fig. 3e**). These data indicate that the loss of ribosome rescue factors in human stem cells results in a modest activation of the RSR.

To test whether the impaired cell growth (**Fig. 3b**) and cytotoxicity (**Fig. 4b**) of *HBS1L* or *ZNF598* loss in hiPSCi can be attributed to activation of the ISR or the RSR, we transduced cells with sgRNAs targeting each of these genes, and induced KRAB-dCas9 expression in the presence of chemical inhibitors of GCN2, PERK, p38, as well as ISRIB, which dampens the ISR^82^ though only at intermediate activation levels^78,83^. Inhibition of GCN2, PERK, or the ISR downstream of eIF2α phosphorylation by ISRIB alleviated the growth and survival defects of hiPSCi depleted for ZNF598 or HBS1L without affecting control cells transduced with a non-targeting sgRNA (**Fig. 4e**, **Extended Data Fig. 3f**). We also analysed the effect of genetically dampening the ISR by co-depleting GCN2 and PERK using dual sgRNA vectors. Knockdown of *GCN2* or *PERK* alleviated the growth and survival defects imparted by ZNF598 or HBS1L depletion in hiPSCi **(Fig. 4f**). Collectively, these data indicate that the loss of ribosome rescue factors in human stem cells impairs cell growth and survival primarily via the ISR.

We next sought to determine the molecular basis for the lack of global translation defects (**Fig. 4a**), cytotoxicity (**Fig. 4b**), and ISR gene expression signatures (**Fig. 4d**) upon ZNF598 or HBS1L depletion in HEK293i. Unlike hiPSCi, HEK293i are aneuploid^19^ and may thus experience constitutive proteotoxic stress due to unbalanced gene expression^84^, making them more resilient to additional perturbations^85^. To test this hypothesis, we compared the levels of total and phosphorylated eIF2α and p38 in hiPSCi and HEK293i by immunoblotting. The levels of phosphorylated eIF2α were ∼40% higher in HEK293i, an effect size comparable to the eIF2α phosphorylation increase triggered by pharmacological induction of ER stress with tunicamycin (**Fig. 4g,h**). The higher levels of eIF2α phosphorylation are in line with the lower *de novo* protein synthesis rates we observed in HEK293i when compared to hiPSCi (**Extended Data Fig. 1g,h**). Moreover, the levels of phosphorylated p38 were ∼100-fold higher in this cell line when compared to hiPSCi (**Fig. 4g,h**). Taken together, these data suggest that constitutive stress signalling may mask the phenotypic consequences of perturbing ribosome rescue in aneuploid cell lines like HEK293.

### Defective ubiquitination by ZNF598 in stem cells elicits ribosome pausing at start sites

The pronounced sensitivity of human stem cells to ZNF598 depletion (**Fig. 2b**, **3b**), suggests that some endogenous mRNAs are difficult to translate and are thus subject to ZNF598-dependent quality control in this cellular context. We reasoned that these quality control events may be difficult to detect in wild-type cells, since ZNF598-mediated r-protein ubiquitylation likely occurs on a millisecond scale^86^ and is followed by rapid ribosome disassembly^61^. They may also be difficult to detect after ZNF598 depletion, since the knockdown of diverse ribosome rescue and release factors commonly activated the ISR in human stem cells (**Fig. 4c,d**)^87–89^. The resulting highly similar gene expression signatures (**Fig. 4c**) and the global inhibition of translation initiation due to eIF2α phosphorylation (**Extended Data Fig. 3d**) may mask the ribosome stalling or collision events on endogenous mRNAs that need to be rescued by each pathway (**Fig. 5a**). Indeed, loss of Hel2, the yeast homolog of ZNF598, also triggers the ISR and concomitantly decreases ribosome collision frequency^90,91^.

**Fig. 5:**
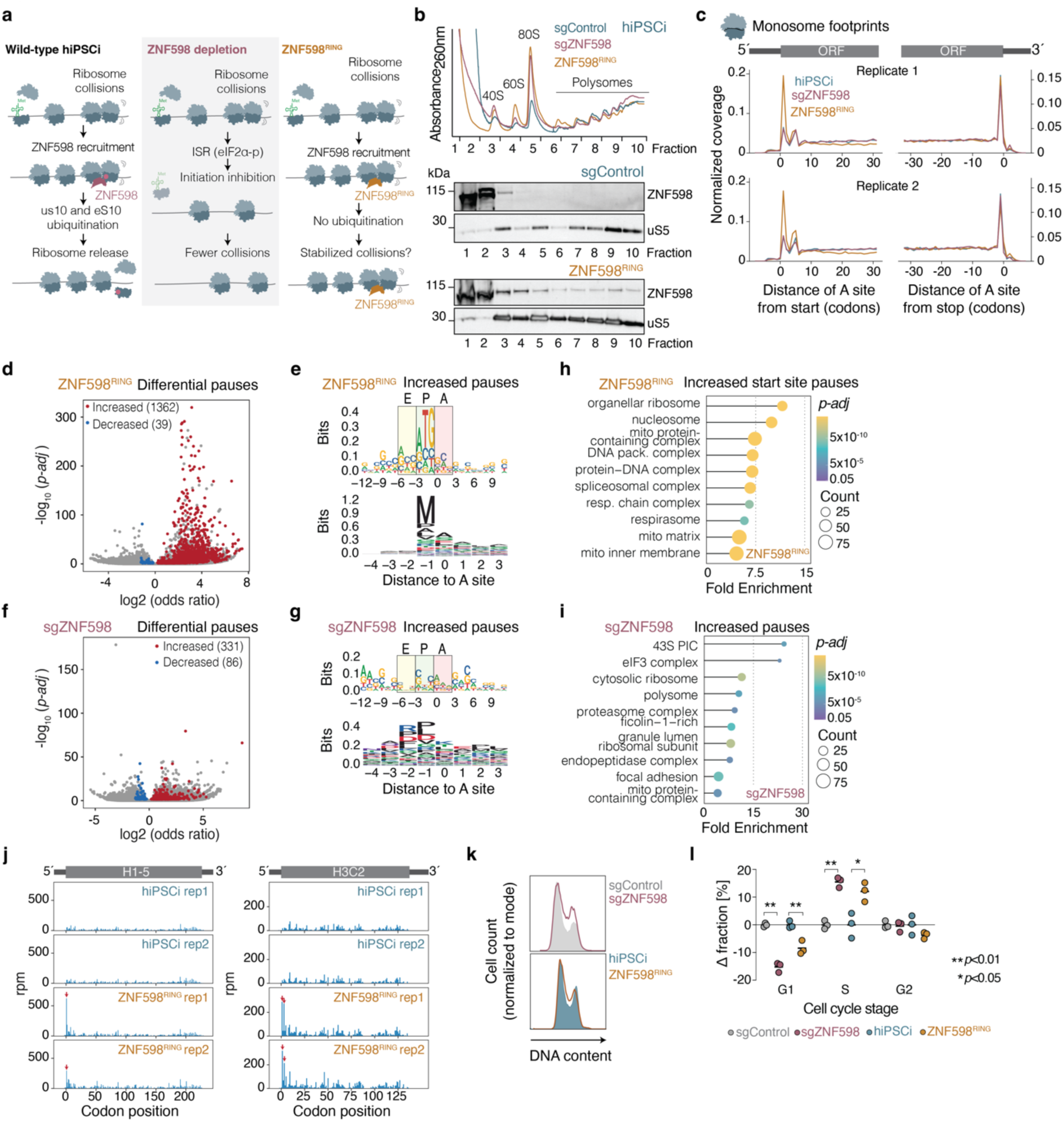
Defective ubiquitination by ZNF598 in human stem cells elicits ribosome pausing at start sites. **a,** Schematic model of the consequences of ZNF598 depletion or ZNF598 RING domain mutant (ZNF598^RING^) expression in hiPSCi. **b,** Polysome profiles (top) and immunoblot analysis of ZNF598 and uS5 in polysome gradient fractions (bottom) of hiPSCi expressing a non-targeting sgRNA (sgControl), a *ZNF598* sgRNA, or ZNF598^RING^. **c,** Metagene profiles of ribosomal A site occupancy from monosome footprints around CDS start and stop sites (*n*=2 biological replicates). **d,** Volcano plot of differential ribosome pause sites upon ZNF598^RING^ expression in hiPSCi (Fisher’s exact-test with Benjamini-Hochberg correction). **e,** Nucleotide (top) and amino acid (bottom) motif analysis of significantly increased pause sites in well-translated mRNAs (> 0.5 footprints/codon in all samples, *n*=3421) in ZNF598^RING^-expressing hiPSCi. **f,** Volcano plot of differential ribosome pausing analysis upon *ZNF598* knockdown (sgZNF598) in hiPSCi as in (**d**). **g,** Nucleotide (top) and amino acid (bottom) motif analysis of significantly increased pause sites in well-translated mRNAs (> 0.5 footprints/codon in all samples, *n*=2463) in sgZNF598 hiPSCi. **h,i**, GO term enrichment analysis of genes with significantly increased pause sites (Fisher’s exact-test with Benjamini-Hochberg correction, *p-adj* ≤ 0.05) within the first five codons in ZNF598^RING^-expressing hiPSCi (**h**) and throughout the ORF in sgZNF598 hiPSCi (**i**) filtered for TPM >1 in RNA-Seq from hiPSCi. **j**, Distribution of monosome footprints (in reads per million, rpm) along the H1-5 (left) and H3C2 (right) mRNA in control and ZNF598^RING^-expressing hiPSCi (*n=*2 biological replicates). Significant differential pauses are indicated with red arrows. **k,** Representative histograms of cell cycle analysis in hiPSCi by DNA staining with EdU followed by flow cytometry. **l,** Changes in the fraction of cells in different cell cycle phases calculated by flow cytometry analysis after EdU staining (*n*=3 biological replicates, >10,000 cells/ analysis; *p* values from unpaired two-tailed t-test).

We postulated that a catalytically inactive ZNF598 would stabilize ribosomes engaged in problematic translation events on endogenous mRNAs. To test this, we substituted the conserved cysteines in the N-terminal RING domain of ZNF598, which should not impair substrate recognition, but block the recruitment and activation of an ubiquitin-conjugating E2^92^. Accordingly, truncated ZNF598 lacking the RING domain can still bind ribosomes, and Cys29 and/or Cys32 substitutions ablate uS10 and eS10 ubiquitination and enhance reporter stalling readthrough even in the presence of endogenous ZNF598 in HEK293T cells^57,58,62^. Overexpression of a Cys29Ser/Cys32Ser RING domain mutant (ZNF598^RING^) but not wild-type ZNF598 increased the expression of mOrange downstream of the AAA_20_ stalling sequence in hiPSCi (**Extended Data Fig. 4a,b**), confirming a dominant-negative role for ZNF598^RING^ also in human stem cells. Unlike the endogenous protein, ZNF598^RING^ indeed robustly co-sedimented with hiPSCi polysomes (**Fig. 5b**), in line with prior observations of more stable association of ZNF598 and Hel2 RING mutants in HEK293 and yeast cells, respectively^58,93^. These data suggest that ZNF598 dissociates less efficiently from its substrates in the absence of r-protein ubiquitination.

Similarly to *ZNF598* knockdown, ZNF598^RING^ expression in hiPSCi decreased *de novo* protein synthesis rates (**Fig. 5b, Extended Data Fig. 4c**) and resulted in significantly elevated levels of LDH release (**Extended Data Fig. 4d),** indicating that defective ZNF598 ubiquitination is cytotoxic in this cellular context. The gene expression changes induced by ZNF598^RING^ in hiPSCi, however, were distinct from those triggered by ZNF598 depletion. Although the levels of more than 1500 mRNAs were significantly altered, ATF4 target genes were not among them (**Extended Data Fig. 4e,f**). eIF2α phosphorylation levels were also not elevated in ZNF598^RING^-expressing hiPSCi, while p38 phosphorylation was only mildly increased (**Extended Data Fig. 4g**). Genes upregulated in ZNF598^RING^-expressing hiPSCi were enriched for GO terms related to metabolism, nucleosome assembly and organization, and RNA processing, while downregulated genes were related to cell growth and development (**Extended Data Fig. 4h**). Taken together, these data indicate that ZNF598^RING^ expression does not activate the ISR in human stem cells, which could result from the inability of eIF2α kinases to recognize the ZNF598-bound conformation of stalled ribosomes^89,91^.

To determine how ZNF598 perturbations impact translation, we analysed ribosome occupancy by high-throughput sequencing of 20-32 nt ribosome-protected mRNA footprint libraries in hiPSCi and HEK293i depleted for ZNF598 or expressing ZNF598^RING^. Because of the high basal levels of ZNF598 in HEK293i (**Extended Data Fig. 2e,f**), we expressed ZNF598^RING^ in cells depleted for endogenous ZNF598 by CRISPRi. Motif analysis of internal ribosomal pauses (*Z* score ≥ 5)^94^, excluding the first 5 and last 5 codons of each CDS) in these monosome footprint libraries revealed the typical enrichment of proline-rich stretches across all conditions^95^ (**Extended Data Fig. 5a,b**). These data confirm our ability to capture known instances of slow elongation, and indicate that neither ZNF598 depletion nor ZNF598^RING^ expression globally alter elongation kinetics in human cells.

We observed a striking increase in monosome footprint density close to annotated CDS start sites upon ZNF598^RING^ overexpression. This increase was not detectable upon ZNF598 depletion (**Fig. 5c**) or wild-type ZNF598 overexpression (**Extended Data Fig. 5c**) in hiPSC, or upon ZNF598 knockdown or ZNF598^RING^ overexpression in HEK293i (**Extended Data Fig. 5d**). In hiPSCi, ZNF598^RING^ expression significantly increased ribosome pausing at 1362 codons in 704 mRNAs (**Fig. 5d**), with a strong enrichment for AUG and methionine in the ribosomal P-site (**Fig. 5e**). The majority of these increased pauses (796; 58%) were within the first 5 codons of the respective CDS (hereafter start site pauses), and had mammalian Kozak consensus-like motifs (RCC<u>AUG</u>G)^96^ (**Extended Data Fig. 5e**). The remaining 566 increased internal pauses had the typical proline-rich internal pausing motif and no distinct nucleotide patterns (**Extended Data Fig. 5f**). By contrast, *ZNF598* knockdown in hiPSCi resulted in only 331 codons with significantly increased pausing, which had nucleotide and amino acid motifs similar to the ones of internal pauses in wild-type hiPSC (**Fig. 5f,g**). The differential pauses elicited by ZNF598 depletion or ZNF598^RING^ expression in HEK293i were also not enriched for specific motifs (**Extended Data Fig. 5g,h**). These data suggest that in human stem cells, ribosomes pause during initiation or in the early stages of elongation when ZNF598-mediated ubiquitination is perturbed by RING domain mutations. The absence of these pauses upon ZNF598 depletion may be due to the global inhibition of translation initiation resulting from increased eIF2α phosphorylation (**Fig. 4e**).

The 704 transcripts with start site pauses in ZNF598^RING^-expressing cells were strongly enriched for GO terms related to DNA packaging and mitochondrial proteins (**Fig. 5h**), in contrast to a modest enrichment of GO terms related to translation among the 313 mRNAs with increased pausing upon ZNF598 depletion (**Fig. 5i**). Forty-one histone-encoding mRNAs had significantly increased ribosome pauses at or shortly after their annotated start sites upon ZNF598^RING^ expression, with little difference in footprint coverage downstream (**Fig. 5j**). The abundance of many of these histone mRNAs and other transcripts with start site pauses in hiPSCi were significantly lower in HEK293i (**Extended Data Fig. 5i,j**), in line with the milder growth defects upon ZNF598 depletion in this cellular context (**Fig. 3b**).

We reasoned that perturbed translation of histone mRNAs could be particularly problematic for stem cells, which progress more rapidly through the cell cycle and have a shortened G1 phase in comparison with somatic cells^97^. Canonical histone-encoding mRNAs are produced exclusively during S phase^98^ and insufficient histones for packaging newly made DNA trigger S-phase arrest in human cells^99^. Indeed, both *ZNF598* knockdown and ZNF598^RING^ overexpression, but not wild-type ZNF598 overexpression, significantly increased the fraction of hiPSCi in S phase at the expense of G1 (**Fig. 5k,l; Extended Data Fig. 5k**). These data indicate that ZNF598-dependent ubiquitination is required for S phase progression in human stem cells.

### ZNF598 detects ribosome collisions during translation initiation

Given that mRNAs with increased start site pauses in ZNF598^RING^-expressing hiPSCi lacked specific nucleotide or amino acid motifs at the stall site apart from a P-site methionine (**Extended Data Fig. 5e**), we examined other potentially distinguishing features. We found that these mRNAs have significantly shorter 5’ UTRs and higher abundance in comparison to other expressed mRNAs in hiPSCi (*p*<0.01, **Fig. 6a**). However, start site pauses did not increase on the majority (87%) of human mRNAs with 5’ terminal oligopolypyrimidine tracts (5’ TOP)^100,101^, which have extremely short 5’ UTRs^102–104^ (**Extended Data Fig. 6a**), suggesting that 5’ UTR length is not the sole determinant of increased ribosome pausing. Since ZNF598 mediates the specific ubiquitination of uS10 and eS10 residues at the 40S-40S interface of collided ribosomes^58,59,73^, we hypothesized that the start site pauses we observed in ZNF598^RING^-expressing hiPSCi could be due to a scanning 43S pre-initiation complex catching up with an initiating 80S ribosome on these messages. This is plausible based on the known time scales of these events: 43S recruitment (∼10 seconds) and 60S joining (∼30 seconds) on model substrates occur on a similar time scale as the transition of initiating ribosomes to elongation (∼30 seconds), whereas the 43S scans the 5’UTR at ∼100 nt per second^105,106^. The potential for start site collisions would be particularly high for messages that are highly efficient in ribosome recruitment. In line with this hypothesis, recent high-throughput measurements of ribosome recruitment to thousands of human 5’ UTRs revealed a 250-fold range in their translational output in human cell extracts^107^. When comparing ribosome recruitment scores for 5’ UTRs determined in this study, we found that they were indeed significantly higher for mRNAs with start site pauses, and in particular for histone mRNAs, than for transcripts without such pauses in ZNF598^RING^-expressing hiPSCi (**Extended Data Fig. 6b**). These data suggest that start site pauses due to defective ubiquitination by ZNF598 occur on messages with highly efficient translation initiation.

**Fig. 6:**
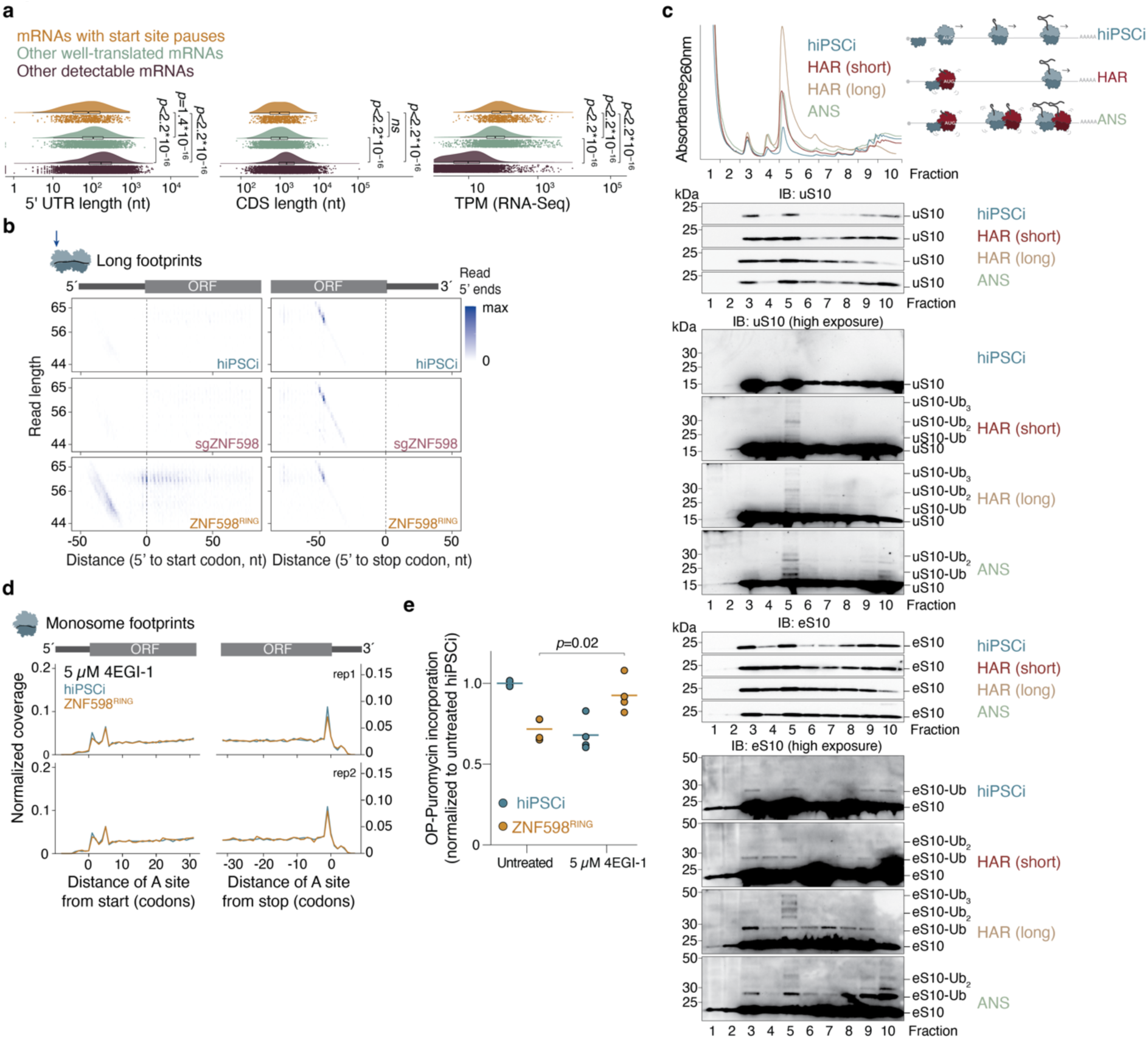
ZNF598 detects ribosome collisions during translation initiation. **a,** Comparison of 5’ UTR length, CDS length, and transcript abundance (by TPM in RNA-Seq) in mRNAs with significantly increased start site pauses (in the first 5 codons of the respective CDS) in ZNF598^RING^-expressing hiPSCi (filtered for TPM>1, *n=*702), other well-translated mRNAs included in the pause site analysis (>0.5 footprints/codon, filtered for TPM>1, *n=*2678), and remaining mRNAs with detectable expression in hiPSCi (TPM>1 in RNA-Seq, *n=*10917). *P*-values from a Wilcoxon test; ns: not significant (*p*>0.01). **b,** Representative density heatmaps of 42-68 nt footprints according to length and 5’ end position around CDS start (left) and stop (right) codons in control, *ZNF598* knockdown (sgZNF598), and ZNF598^RING^-expressing hiPSCi (top to bottom) in one of two biological replicates. **c,** Polysome profiling (top) and immunoblot analysis of sucrose gradient fractions (bottom) from untreated hiPSCi and after treatment with 0.05 mg/l anisomycin (ANS) for 15 minutes, or a short (2.5 min) or long (2 hours) treatment with 2 µg/ml homoharringtonine (HAR). **d,** Metagene profiles of ribosomal A sites from monosome footprints around CDS start and stop codons after treatment with 5 µM 4EGI-1 (*n=*2 biological replicates). **e,** Global protein synthesis measurements by O-propargyl-puromycin (OPP) labeling in control or ZNF598^RING^-expressing hiPSCi before and after treatment with 5 µM 4EGI-1. Median fluorescence intensity quantified by flow cytometry (>10,000 cells/ analysis) was normalized to values from untreated hiPSCi (*n=*3-4 biological replicates).

We next asked whether we can detect 40S-80S collisions near translation start sites by ribosome profiling. Capturing translation steps prior to 80S assembly typically requires protein-RNA crosslinking^108^, but we reasoned that ZNF598^RING^ may stabilize 40S-80S collisions and render their footprints detectable without this additional step. Mammalian 43S complexes protect fragments of 20 to 60 nt from nuclease digestion^109,110^, while 80S di-ribosomes yield ∼60-nt footprints^111,112^. We therefore analysed 50-80 nt (long) ribosome footprints from control hiPSCi and upon ZNF598^RING^ expression or *ZNF598* knockdown. We detected ∼60-nt footprints with 5’ ends mapping ∼50 nt from annotated stop codons in all conditions (**Fig. 6B** and **S6C**), consistent with 80S ribosome collisions during translation termination^90,112^ that are not substrates for ZNF598. A striking additional density of footprints 45 to 62 nt in length was detectable only in ZNF598^RING^-expressing hiPSCi. The 5’ ends of these footprints mapped 20 to 40 nt upstream of annotated start codons, and their 3’ ends mapped ∼18 nt downstream. Footprints of ∼60 nt length with 5’ ends around annotated start sites, indicative of 80S-80S collisions during early elongation, were also detectable only in ZNF598^RING^-expressing hiPSCi (**Fig. 6b; Extended Data Fig 6c,d**). Metagene analysis confirmed an accumulation of long footprints with P sites at or immediately after start codons, as well as ∼30 nt (corresponding to the footprint of a single ribosome) downstream (**Extended Data Fig. 6e**). Together, these data suggest that ZNF598^RING^ expression in hiPSCi stabilizes 40S-80S collisions during translation initiation and 80S-80S collisions during early rounds of elongation.

We next sought to determine whether increasing the potential for ribosome collisions during initiation would lead to ZNF598-mediated ubiquitination of uS10 and eS10. To test this, we exposed hiPSCi to two translation inhibitors: homoharringtonine, which blocks the first peptide bond formation^113^, and anisomycin, which at intermediate concentrations induces collisions of elongating ribosomes that trigger p38 phosphorylation^55^. We reasoned that both drugs could result in 40S-80S collisions as they inhibit translation elongation but do not inhibit 40S loading onto mRNA; detection of these collisions by ZNF598 should result in uS10 and eS10 ubiquitination in monosome fractions. Brief (2.5 min) or extended (2 hours) homoharringtonine treatment of hiPSCi led to polysomal collapse and a concurrent increase in the 80S peak, indicating efficient ribosome run-off (**Fig. 6c**). In line with our predictions, we detected uS10 polyubiquitination in 80S fractions from hiPSCi treated with both inhibitors, as well as in heavy polysome fractions from cells treated with anisomycin under conditions that induce p38 phosphorylation (**Fig. 6c**, **4g**). Monoubiquitinated eS10, which is not sufficient to trigger ribosome subunit dissociation^60^, was present in 40S, 80S, and heavy polysome fractions in untreated hiPSCi. However, eS10 poylubiquitination increased specifically in 80S fractions upon treatment with both drugs, and also in heavy polysome fractions upon anisomycin exposure (**Fig. 6c**). In HEK293i, by contrast, a 2.5-min homoharringtonine treatment did not lead to uS10 ubiquitination and only mildly increased eS10 monoubiquitination (**Extended Data Fig. 6f**), consistent with the higher basal levels of eIF2α phosphorylation in these cells (**Fig. 4g**) that likely limits 40S loading. Collectively, these data suggest that ZNF598 ubiquitinates uS10 and eS10 on ribosomes colliding at translation start sites in human stem cells. This pathway, however, is distinct from the initiation RQC (iRQC), in which the E3 ligase RNF10 ubiquitinates uS3 and uS5 to flag ribosomes terminally stalled during initiation due to 40S rRNA defects or prolonged homoharringtonine exposure^114–117^. Consistently, uS3 and uS5 ubiquitination increased substantially in hiPSCi upon prolonged (2 hours) but not brief (2.5 min) homoharringtonine treatment (**Extended Data Fig. 6g**), suggesting that ZNF598 detects start-site ribosome collisions in early or mild stress conditions before the iRQC is activated. If ZNF598 detects a scanning 43S collided with an 80S ribosome that is transitioning to elongation, reducing 43S loading on mRNAs should decrease the potential for such collisions. To test this prediction, we treated hiPSCi with 4EGI-1, an inhibitor of eIF4E/eIF4G complex formation^118^, which we used at a concentration that induces only minimal (∼25%) reduction of global protein synthesis (**Fig. 6d; Extended Data Fig. 6h**). Strikingly, this mild 4EGI-1 treatment was sufficient to abolish the excess ribosome density at start sites and rescue the global translation defects of ZNF598^RING^-expressing cells (**Fig. 6d,e**). Taken together, our data are consistent with a model in which ZNF598 detects collisions of scanning 43S with 80S ribosomes at start codons on mRNAs with high initiation efficiency (**Fig. 7**).

**Fig. 7:**
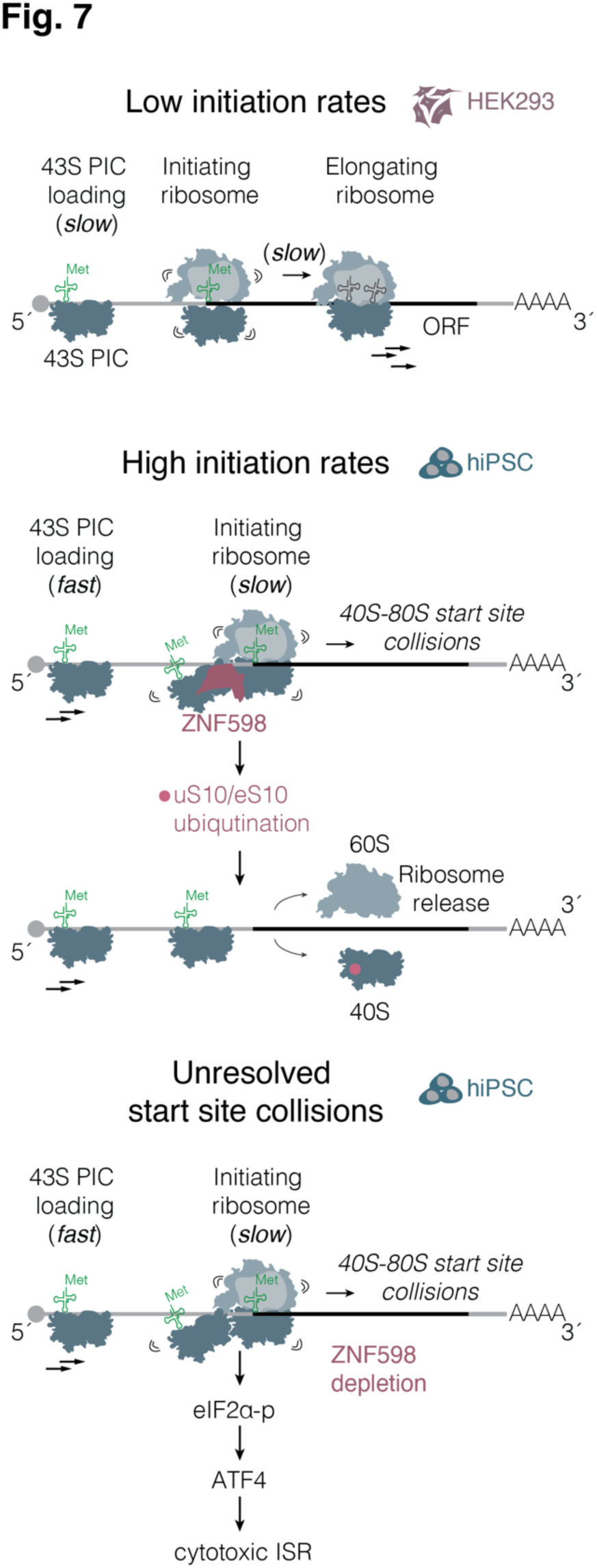
Model for ZNF598-mediated surveillance of ribosome collisions during translation initiation. PIC: pre-initiation complex; ORF: open reading frame.

## Discussion

By mapping the essentiality of mRNA translation-related genes across diverse hiPSC-derived cell types, our study provides a rich resource for dissecting pathway-specific regulatory mechanisms in physiologically relevant settings. Through optimized CRISPRi screening and differentiation workflows, we achieved robust and efficient gene repression not only in dividing human cells (hiPSC, NPC), but also in post-mitotic hiPSC-derived neurons and cardiac cells. This powerful comparative approach enabled us to identify core and cell type-specific genetic dependencies. In particular, we discovered that human stem cells express mRNAs that display highly efficient translation initiation and require the ribosome collision sensor ZNF598 to prevent activation of a cytotoxic ISR.

The differential essentiality of translation-coupled quality control pathways we identified here suggests that their endogenous triggers are limited to defined cell types or cell-fate transitions. The consistent patterns we observed for genes encoding subunits of a protein complex (e.g., *GIGYF2* and *EIF4E2*, *DNAJC2* and *HSPA14*) lend further support to this idea. In some cases, we uncovered a discordant essentiality among genes that are thought to work in the same pathway, pointing to potential redundancy or independent, specialized roles for individual pathway components (e.g., *EDF1* and *ZNF598; PELO* and *HBS1L*) in distinct cellular contexts.

Our finding that *ZNF598* or *HBS1L* repression triggers a cytotoxic ISR rationalizes the essentiality of these genes in human stem cells. Activation of the ISR, which also happens upon deletion of the *ZNF598* homolog *HEL2* in yeast, may decrease ribosomal flux along messages, limiting the chances for ribosome stalling and collisions^90^. Along with the extensive reprogramming of gene expression that accompanies the ISR, decreased ribosome flux may however mask the endogenous translation events that require surveillance, explaining why their discovery has been so challenging. The common activation of the ISR upon deletion or repression of distinct ribosome quality control genes^87–89^, but not upon expression of their corresponding catalytically inactive mutants^91^, suggests that the broader use of such mutants may be better suited for identifying the physiological triggers of these pathways in cells. Our data also indicate that a basal level of ISR activation in HEK293, and potentially other immortalized cell lines, could contribute to their resilience to perturbations in ribosome rescue^57^, further confounding its mechanistic dissection.

We find that human stem cells are especially reliant on the pathways that detect and rescue slow or stalled translating ribosomes. These pathways were initially discovered in yeast with the help of mRNA reporters containing unnaturally strong RNA hairpins or rare codon stretches^49,119^. The most widely used sequences in mammalian stalling reporters are the XBP1 arrest peptide, or poly(A) repeats of more than 36 nucleotides^58,73^. Poly(A) stretches, however, trigger ribosome sliding and frame loss^120^ and are strongly underrepresented in human genes^121^. Ribosomes could in theory encounter poly(A) repeats when translating a prematurely polyadenylated mRNA^57,73^, but eukaryotes have dedicated mechanisms that suppress premature mRNA cleavage and polyadenylation^122^. Apart from the XBP1 arrest peptide, there are only a handful of other examples of nascent chain sequences that stall mammalian ribosomes. These sequences are mostly encoded by upstream ORFs^123^, which would obviate the need for peptide elongation past the stall. This underrepresentation of ribosome stalling sequences in mammalian genes has been a barrier to deciphering the physiological roles of ribosome rescue pathways, and our data also indicate that these pathways are required only in some cellular contexts.

By identifying the essential role of ZNF598 in human stem cells and profiling the consequences of its catalytic inactivation, which unlike ZNF598 depletion does not induce the ISR and therefore does not inhibit ribosome loading, we discovered a previously unappreciated source of problematic translation events in stem cells: the slow transition of initiating ribosomes to elongation. This transition is two orders of magnitude slower than subsequent elongation rounds^105,106^, and our data suggest that ZNF598 detects queuing and collisions that occur if a scanning 43S catches up with an initiating 80S. Structures of ZNF598 are still lacking, but uS10 and eS10, which are targeted for ubiquitination by ZNF598, reside at the 40S-40S interface in collided di-ribosomes^59,60,124^. The absence of 60S contacts in these structures supports the notion that the surface recognized by ZNF598 could be formed also by a 40S-80S collision. Mechanisms to detect and resolve such collisions would ensure the accuracy and fidelity of start site selection by preventing frameshifting of the initiating 80S^125^. Our data indicate that such collisions do not occur on all mRNAs or in all cell contexts: in HEK293 cells, uS10 was not ubiquitinated upon ribosome stalling at CDS start sites induced by a brief exposure to homoharringtonine, in line with the absence of 40S queued at start sites in similarly treated HeLa cells^109^. This could be due to the higher basal levels of eIF2α phosphorylation upon persistent ISR activation in immortalized cell lines like HEK293 and HeLa that globally limits ribosome loading on messages, or to differences in the stoichiometry of translation initiation factors among cell types.

We find that start site collisions occur in a subset of mRNAs with initiation sites in a strong Kozak context and with shorter than average 5’ UTRs that need to be translated at high levels and may thus have particularly high ribosomal loading rates^107^. Poignant examples of such transcripts are those encoding histone proteins, which are synthesized in large amounts only during S phase. Interestingly, ribosomes are loaded on newly made histone mRNAs approximately 5 times more rapidly than on housekeeping mRNAs also in mouse embryonic stem cells^126^. Such rapid loading would enable cells to quickly produce large amounts of protein from these messages, but may also increase the probability of 40S-80S collisions at start sites. The larger fraction of stem cells in S phase^97^ and their broader repertoire of expressed histone genes may therefore partially account for their selective reliance on ZNF598.

Interestingly, we did not detect start site pauses in most 5’ TOP mRNAs, which have extremely short 5’ UTRs that may not be compatible with the recruitment of a 43S before initiating ribosomes have cleared the AUG^127^. Extremely short 5’ UTRs could thus have evolved to protect messages from start site ribosome collisions, although this would need to be carefully balanced with the increased chances of leaky scanning^128^. The median length of yeast 5’ UTRs is only 50 nt^129^, in contrast to the ∼200-nt median 5’ UTR length in human mRNAs. This difference could account for the lack of ribosome accumulation at start sites in yeast cells expressing a Hel2 RING domain mutant^90^. It is conceivable that Hel2 and ZNF598 have distinct endogenous substrates, given that they share only 15% sequence identity^57^ and that their main targets on the 40S subunit also differ^73,130^. Our discovery of ZNF598’s role in detecting ribosome collisions during translation initiation in human stem cells emphasizes the need to study ribosome quality control pathways in biological contexts where they are necessary to maintain the translational homeostasis of endogenous mRNAs.

## Acknowledgments

We thank K. Strasser for expert technical assistance, the NGS Facility in the Department of Totipotency at MPIB for high-throughput sequencing, the Mass Spectrometry Core Facility at MPIB for proteomics measurements, the Imaging Core Facility at MPIB for help with FACS measurements, the Protein Production Core Facility at MPIB for recombinant TS2126 RNA ligase 1, and N. Sinha and K. Stein for advice. G.R., A.B. and S.W. were supported by the International Max Planck Research School for Molecular Life Sciences (IMPRS-LS). S.F. was supported by a postdoctoral fellowship from the Alexander von Humboldt Foundation. This work was funded by the Max Planck Society, the European Research Council under the European Union’s Horizon 2020 Research and Innovation Programme (ERC Starting Grant No. 803825-TransTempoFold to D.D.N.), the EMBO Young Investigator Program (YIP 4833 to D.D.N.) and the Deutsche Forschungsgemeinschaft (DFG, German Research Foundation) through Germany’s Excellence Strategy - EXC 2145 – 390857198 to D.E.).

## Author contributions

Conceptualization, G.R. and D.D.N.; Experimental methodology, G.R., S.W. and D.D.N.; Investigation, G.R.; Formal analysis, G.R., A.B., S.F., S.W., D.D.N; Resources (HEK293i cell line): H.R. and D.E.; Writing - original draft, G.R. and D.D.N.; Writing – review & editing, G.R., S.F., A.B., S.W., D.D.N.; Supervision and funding acquisition, D.D.N.

---

**Extended Data Fig. 1:**
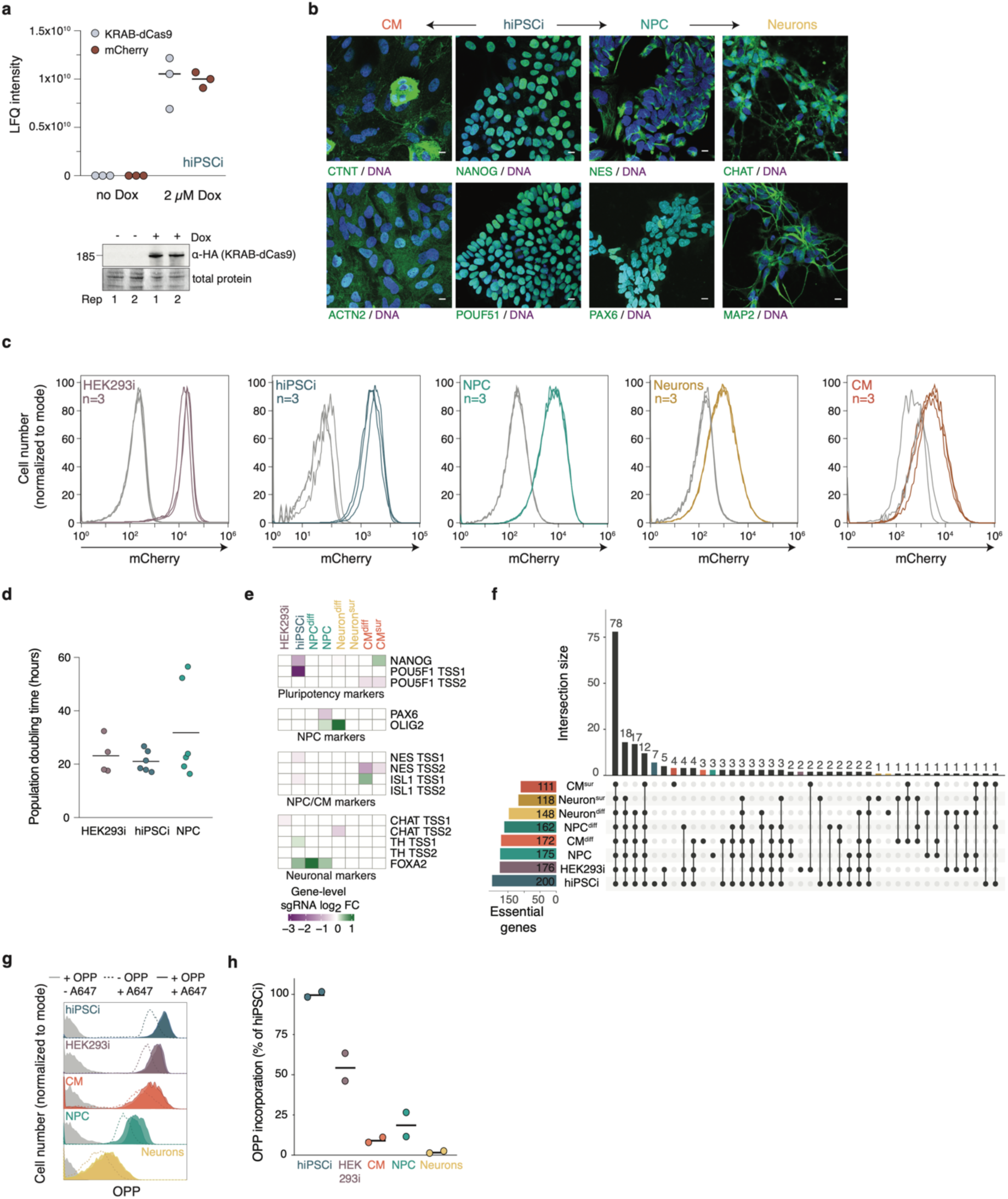
Workflow for comparative genetic screens by inducible CRISPRi in HEK293 and hiPSC-derived cells. **a,** LFQ intensities of KRAB-dCas9 and mCherry protein levels in hiPSCi +/- doxycycline (Dox) measured by mass spectrometry (*n=*3 biological replicates, top) and immunoblot analysis of HA-tagged KRAB-dCas9 in control hiPSCi and after treatment with 2 µM doxycycline for 2 days (bottom). **b,** Immunostaining for cell type-specific markers (green) and DAPI (blue). Scale bar: 10 µm. **c,** Quantification of % mCherry-positive cells by flow cytometry (*n=*3 biological replicates) after KRAB-dCas9 induction with 2 µM doxycycline for two days (in hiPSCi and HEK293i), 5 passages (NPC), and for five days in differentiated neurons and CM. **d,** Quantification of cell doubling time (*n=*2 biological replicates for HEK293i; *n=*3 biological replicates for hiPSCi and NPC). **e,** Heatmaps of significant gene-level sgRNA log_2_ fold change (FC) (*p* ≤ 0.1) for cell identity markers. diff = differentiation, sur = survival, TSS = transcriptional start site. **f,** UpSet plot of overlap among genes with significant negative gene-level sgRNA log_2_ FC (Mann-Whitney *p* ≤ 0.1). **g,h**, Global protein synthesis measurements by O-propargyl-puromycin (OPP) labeling. (**g**) Median fluorescence intensity quantified by flow cytometry (>10,000 cells/ analysis); (**h**) Data from (**e**) normalized to a matched control not treated with OPP (-OPP + A647) and calculated as a fraction of the average signal in hiPSCi (*n*=2 biological replicates for +OPP+A647; *n*=1 biological replicate for -OPP+A647 and +OPP-A647).

**Extended Data Fig. 2:**
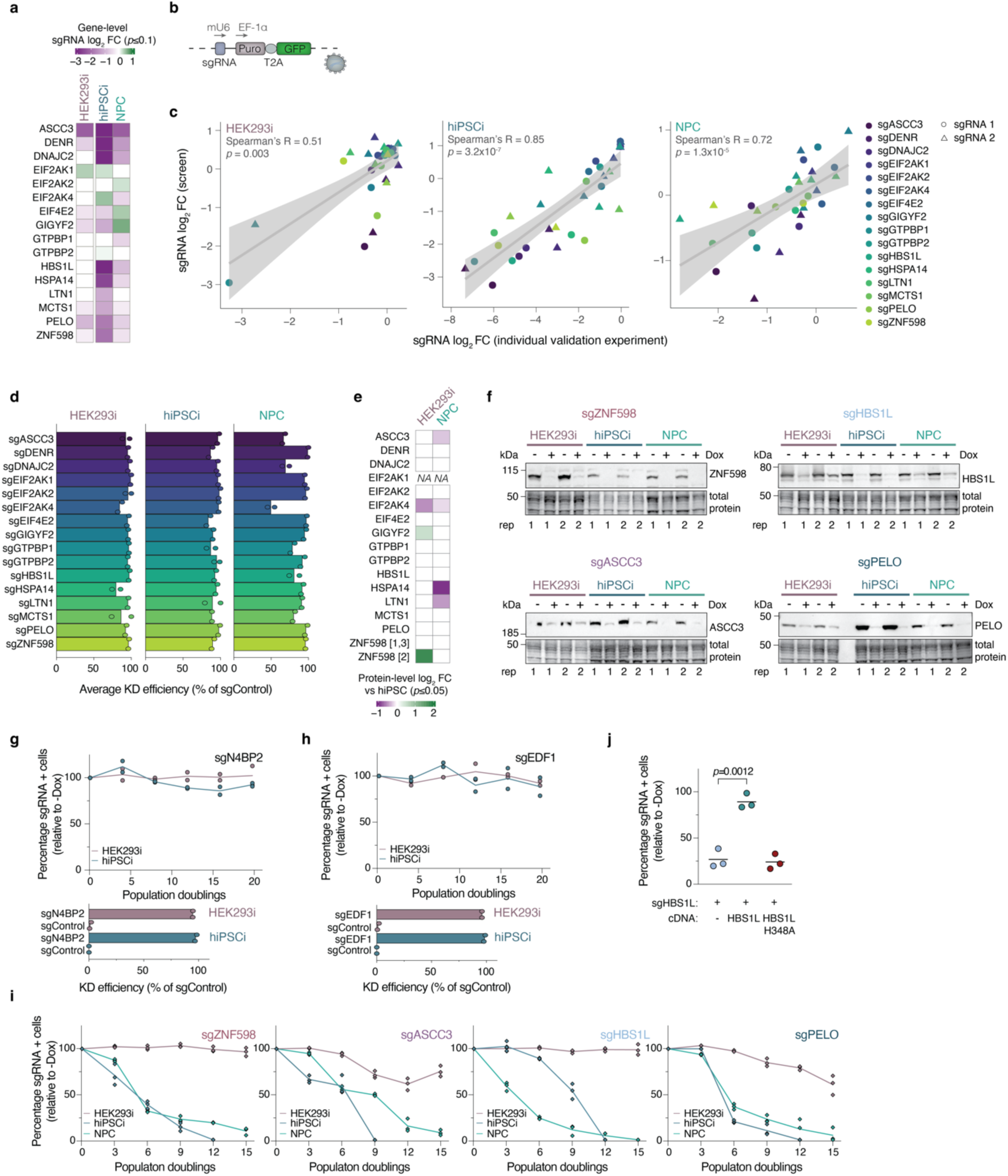
Validation of pooled CRISPRi screens in dividing cells. **a,** Heatmaps of gene-level sgRNA log_2_ FC in hiPSCi, HEK293i, and NPC of 16 genes selected for validation; white: not significant (Mann-Whitney *p* > 0.1). **b,** Schematic of individual sgRNA expression construct. Puro: puromycin resistance gene. **c,** Correlation of single sgRNA phenotype from growth competition assays (log_2_ (-/+Dox)) and log_2_ FC of the same sgRNA in the pooled CRISPRi screen (Spearman’s R, *p*< 0.05) for the two most active sgRNAs (sgRNA1 and sgRNA2) for each gene in the screens. **d,** Knockdown efficiency of genes from (**a**) compared to a non-targeting control (sgControl) measured by quantitative RT-PCR for the most active sgRNA from the screens (sgRNA1; *n=*2 biological replicates, each with *n=*3 technical replicates). **e,** Heatmaps of significant (pairwise two-sided t-test, FDR<0.01) protein-level log_2_ FC for screening targets in (**a**) measured by mass spectrometry (*n=*3 biological replicates). Numbers in brackets indicate differentially detected isoforms. **f**, Immunoblot analysis of ZNF598, ASCC3, HBS1L, and PELO in sgRNA-transduced cells after KRAB-dCas9 induction (+Dox) and in matched uninduced controls (-Dox). **g,h,** Growth assays of HEK293i or hiPSCi expressing a sgRNA targeting N4BP2 (**g**) or EDF1 (**h**) (*n*=2 biological replicates; line: average) and knockdown efficiency measurements of target genes compared to a non-targeting control (sgControl) by quantitative RT-PCR (*n*=2 biological replicates, each with *n*=3 technical replicates). **i,** Growth assays of cells transduced with the second most potent sgRNA (sgRNA2) targeting *ZNF598*, *ASCC3*, *HBS1L*, or *PELO* in the hiPSCi screen (*n=*3 biological replicates; line: average). The percentage of GFP-positive (GFP+) cells was measured by flow cytometry (>10,000 cells/ analysis) every three population doublings and normalized to GFP+ cell numbers in matched uninduced (-Dox) controls. **j,** Growth assays of hiPSCi transduced with a sgRNA targeting *HBS1L* alone or in combination with a cDNA encoding wild-type or a GTPase-inactive *HBS1L* (H348A) (*n*=3 biological replicates, *p*-values from unpaired two-tailed t-test).

**Extended Data Fig. 3:**
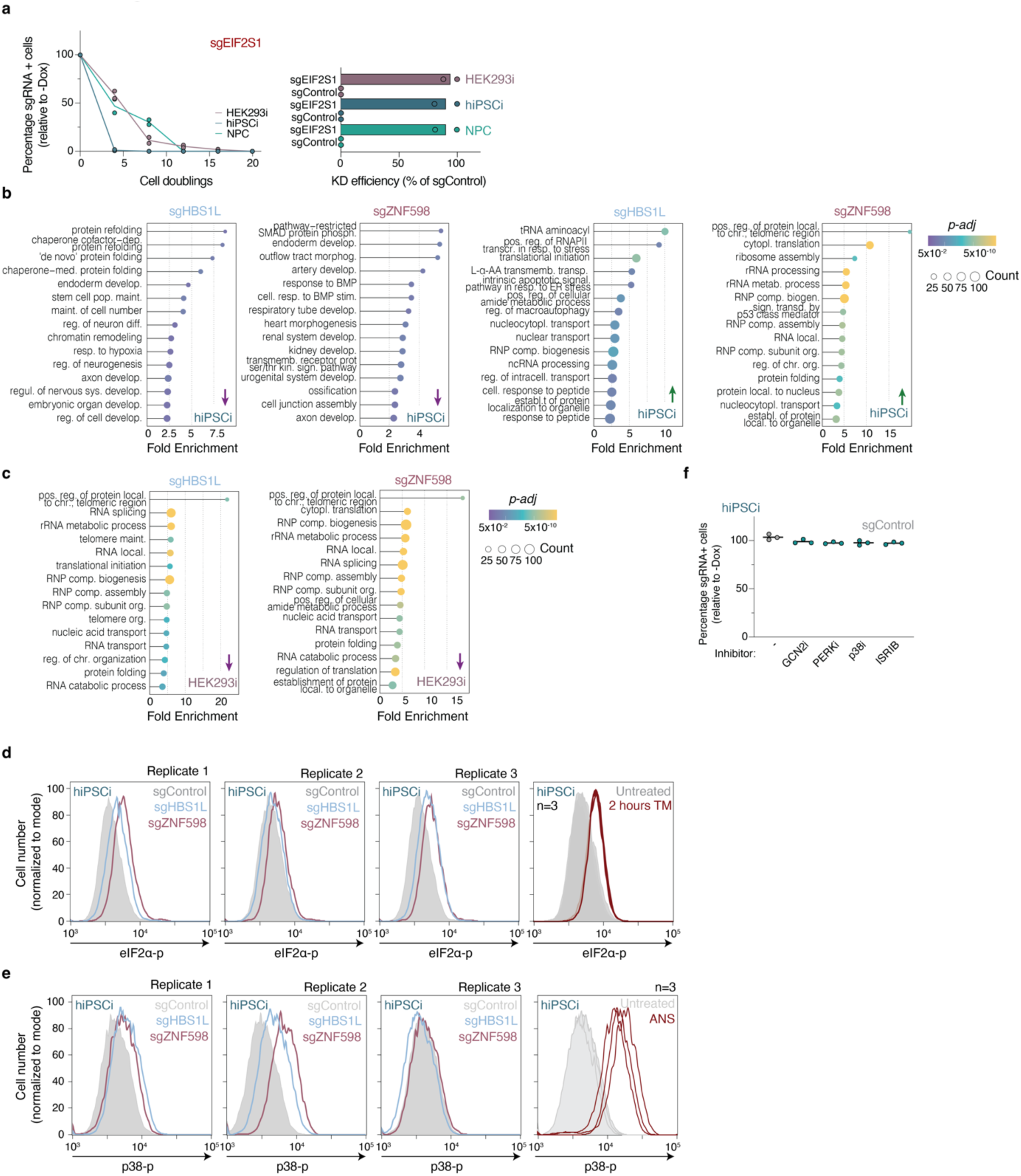
Analysis of stress pathway activation in different cellular contexts. **a,** Growth assays of cells transduced with a sgRNA targeting *EIF2S1* in HEK293i, hiPSCi, or NPC cells (*n=*2 biological replicates; line: average) and knockdown efficiency measurements of target genes compared to a non-targeting control (sgControl) by quantitative RT-PCR (*n*=2 biological replicates, each with *n*=3 technical replicates). **b,c**, Gene ontology (GO) enrichment in differentially expressed mRNAs using the “Biological Process” function in ClusterProfiler (Benjamini-Hochberg FDR <0.05) in knockdown hiPSCi (**b**) or HEK293i (**c**). **d,e**, Flow cytometry analysis of eIF2α phosphorylation (**d**) and p38 phosphorylation (**e**) upon depletion of ZNF598 or HBS1L in hiPSCi in comparison to cells transduced with a non-targeting sgRNA (sgControl; n=3 biological replicates; matched samples). hiPSC treated with 2.5 µg tunicamycin (TM) for 2 hours to induce the ISR or with 0.05 mg/l anisomycin (ANS) for 15 minutes to induce the RSR served as a positive controls. **f**, Growth assays of hiPSCi expressing a non-targeting sgRNA (sgControl, day 6) in the absence (-) or presence of inhibitors of GCN2 (GCN2i, A-92, 1.25 µM), PERK (PERKi, GSK2606414, 4 nM), p38 (p38i, SB203580, 1 µM), or the ISR (ISRIB, 50 nM) in comparison to uninduced controls (*n*=3 biological replicates, *p*-values from unpaired two-tailed t-test).

**Extended Data Fig. 4:**
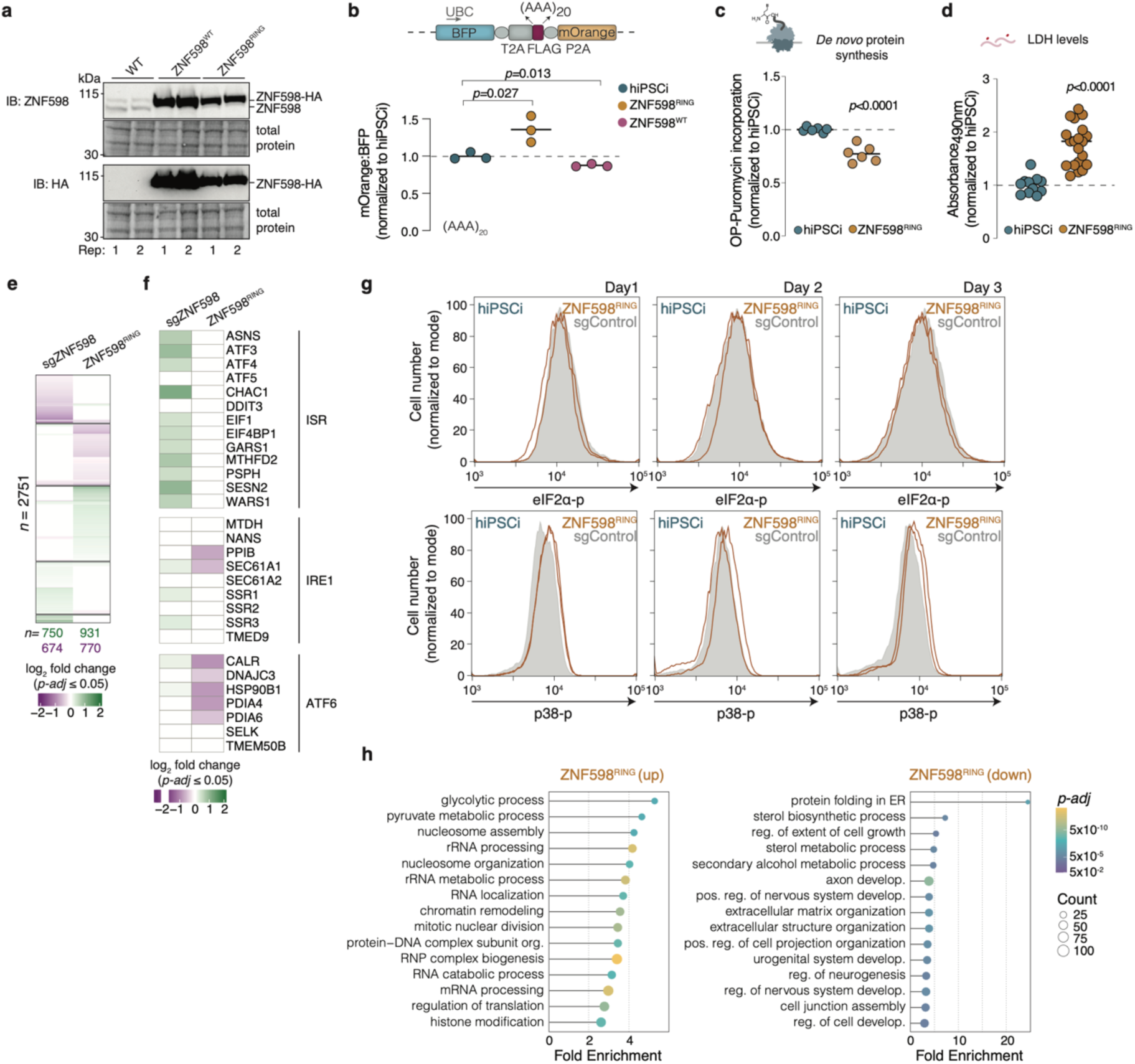
ZNF598^RING^ expression in human stem cells induces cytotoxicity but not through the ISR. **a,** Immunoblot analysis of wild-type hiPSCi and hiPSC transduced with cDNA encoding HA-tagged wild-type (ZNF598^WT^) or C29S/C32S (ZNF598^RING^) *ZNF598* (*n*=2 biological replicates). **b,** Stalling readthrough of reporters containing an AAA-encoded stretch of twenty lysines (AAA_20_) in control hiPSCi and hiPSCi transduced with ZNF598^WT^ or ZNF598^RING^ cDNA. The median fluorescence intensity for BFP and mOrange was quantified by flow cytometry (>20,000 cells/ analysis) and normalized to average values for the hiPSCi control (*n*=3 biological replicates; *p* values from unpaired two-tailed t-test). **c,** Global protein synthesis measurements by OP-Puromycin and AF647 detection. Median fluorescence intensity was quantified by flow cytometry (>10,000 cells/ analysis) and normalized to average values for the control (*n*=2 biological replicates; *p* values from unpaired two-tailed t-test). **d,** Lactate dehydrogenase (LDH) measurements in culture supernatants from wild type or ZNF598^RING^-expressing hiPSCi two days after transduction. Values were normalized to supernatant from wild type hiPSCi (*n=*4 technical replicates and ≥ 3 biological replicates; *p*-values from unpaired two-tailed t-test). **e,f**, Differential gene expression analysis in hiPSCi upon ZNF598 knockdown (sgZNF598) in comparison to a non-targeting sgRNA (sgControl), or in ZNF598^RING^-expressing hiPSCi in comparison to untransduced hiPSCi (*n*=2 biological replicates). (**e**) Heatmap of mRNAs differentially expressed in at least one context (Wald test; *p-adj* ≤ 0.05; *n*=2289). (**f**) Heatmap of a subset from (**e**) showing genes associated with the integrated stress response (ISR). **g,** Flow cytometry analysis of eIF2α (top) or p38 (bottom) phosphorylation upon ZNF598^RING^ expression in hiPSCi in comparison to cells transduced with a non-targeting sgRNA (sgControl; *n*=2 biological replicates) on day 1-3 after transduction. **h,** Gene ontology (GO) enrichment in differentially expressed mRNAs from (**e**) using the “Biological Process” function in ClusterProfiler (Benjamini-Hochberg FDR <0.05).

**Extended Data Fig. 5:**
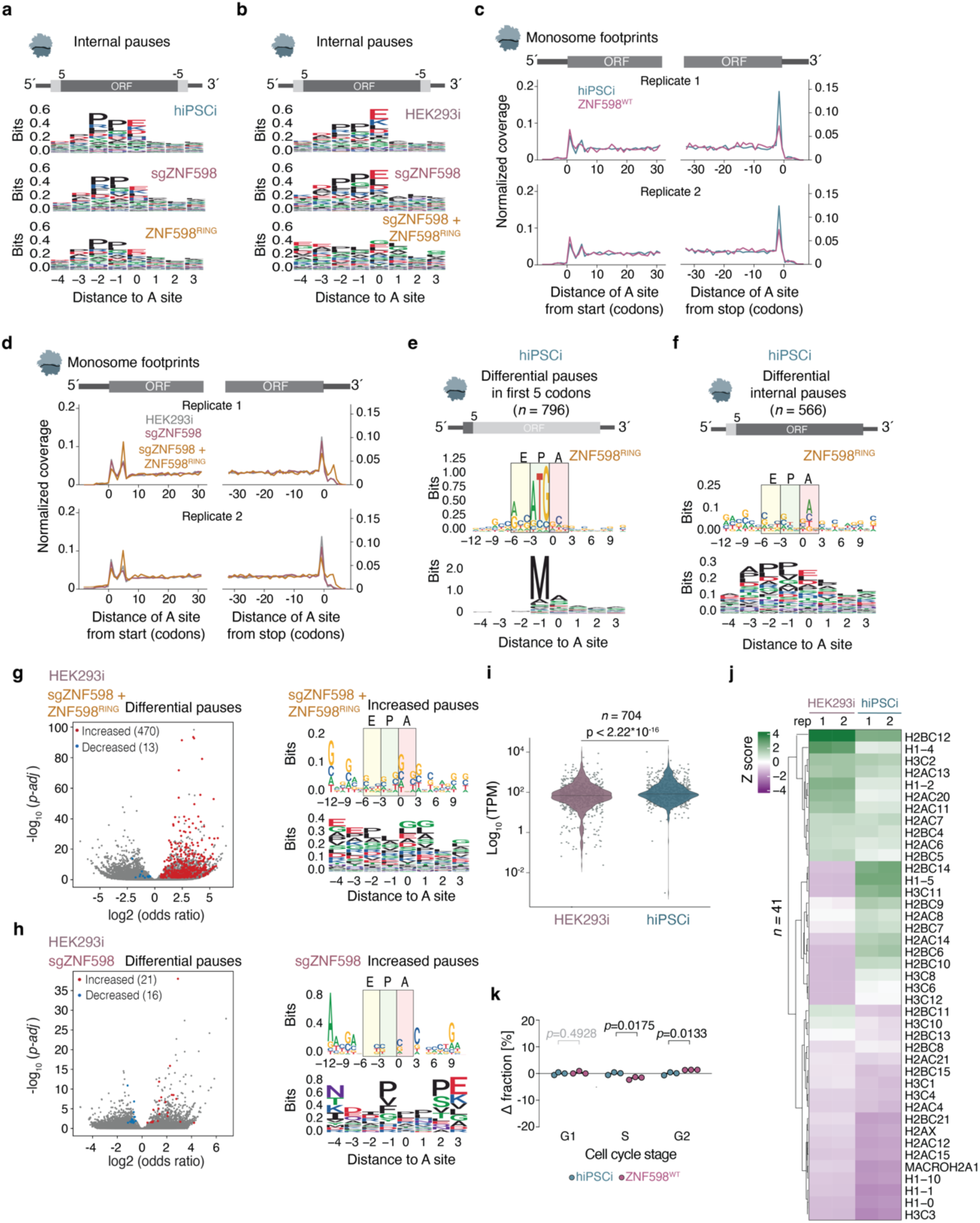
ZNF598^RING^ increases ribosome occupancy at translation start sites in human stem cells. **a,b**, Amino acid motif analysis of internal pause sites (excluding the first and last five codons of each CDS) in wild type, *ZNF598* knockdown (sgZNF598), and ZNF598^RING^-expressing hiPSCi (**a**) or HEK293i (**b**). **c,** Metagene profiles of ribosomal A sites from monosome footprints around CDS start and stop codons in wild type and ZNF598^WT^-overexpressing hiPSCi (*n=*2 biological replicates). **d,** Metagene profiles of ribosomal A sites from monosome footprints around CDS start and stop codons in wild type, sgZNF598 and ZNF598^RING^-expressing HEK293i (*n=*2 biological replicates). **e,f**, Nucleotide (top) and amino acid (bottom) motif analysis of significantly increased pause sites within (**e**) or excluding (**f**) the first five codons in ZNF598^RING^-expressing hiPSCi (Fisher’s exact-test with Benjamini-Hochberg correction, *p-adj* ≤ 0.05). **g,** Volcano plot of differential ribosome pause sites upon ZNF598^RING^ expression in HEK293i (Fisher’s exact-test with Benjamini-Hochberg correction) (left). Nucleotide (top) and amino acid (bottom) motif analysis of significantly increased pause sites in well-translated mRNAs (>0.5 footprints/codon in all samples, *n*=1438) in ZNF598^RING^-expressing HEK293i (right). **h,** Volcano plot of differential ribosome pausing analysis upon *ZNF598* knockdown (sgZNF598) in HEK293i as in (**g**, left). Nucleotide (top) and amino acid (bottom) motif analysis of significantly increased pause sites in well-translated mRNAs (> 0.5 footprints/codon in all samples, *n*=1003) in sgZNF598 HEK293 as in (**g**, right). **i,** Violin plots (center line: median) of TPM values from RNA-Seq for genes with significantly increased A-site pausing within the first 5 codons in ZNF598^RING^-expressing hiPSCi (*n*=704) in sgControl HEK293i and hiPSCi (*p*-value from Wilcoxon test). **j,** Hierarchically clustered heatmap of scaled *Z* scores of normalized unique transcript counts (TPM) from RNA-Seq in sgControl HEK293i and hiPSCi (*n*=2 biological replicates) for histone mRNAs with increased A-site pausing within the first 5 codons in ZNF598^RING^-expressing hiPSCi (*n*=41). **k,** Changes in the fraction of cells in different cell cycle phases calculated by flow cytometry analysis after EdU staining (*n*=3 biological replicates, >10,000 cells/ analysis; *p* values from unpaired two-tailed t-test).

**Extended Data Fig. 6:**
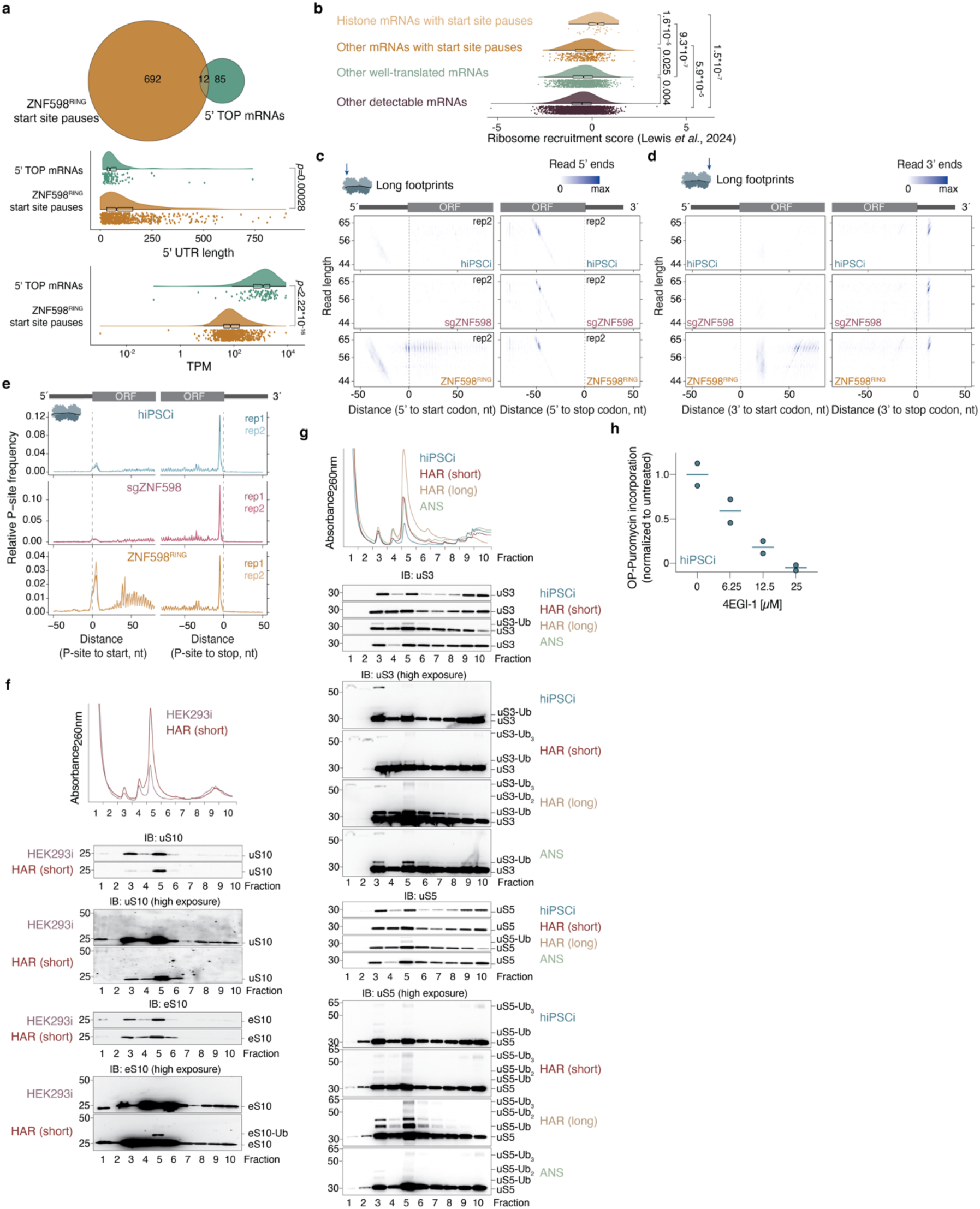
Analysis of long ribosome footprints and cellular responses to initiation site stalling. **a**, Venn diagram of overlaps between 5’TOP motif mRNAs (*n*=97) and transcripts with significantly increased A-site pauses in the first 5 codons in ZNF598^RING^-expressing hiPSCi (*n*=704), and comparison of 5’ UTR lengths and TPM levels from RNA-Seq in both groups (*p*-value from Wilcoxon test). **b,** Comparison of ribosome recruitment scores for 5’UTRs quantified by direct analysis of ribosome targeting (DART)^107^ for histone mRNAs with start site pauses (*n*=29), other mRNAs with start site pauses (*n*=351), other well-translated mRNAs (*n*=1340), and other detectable mRNAs (*n*=3791). **c,** Density heatmaps of 50-80 nt ribosome footprints according to length and 5’ end position around CDS start (left) and stop sites (right) for a second biological replicate. **d,** Density heatmaps of 50-80 nt ribosome footprints according to length and 3’ end position around CDS start (left) and stop sites (right) for a second biological replicate. **e,** Metagene profiles of 50-80 nt ribosome footprints around CDS start and stop sites (*n*=2 biological replicates). **f,** Polysome profiles and immunoblot analysis of uS10 and eS10 in sucrose gradient fractions from control HEK293i or after a short (2.5 min) treatment with 2 µg/ml homoharringtonine (HAR). **g,** Polysome profiles (from Fig. 6c) and immunoblot analysis of uS3 and uS5 in sucrose gradient fractions from control hiPSCi, after short (2.5 min) or long (2 hours) treatment with 2 µg/ml homoharringtonine, or after treatment with 0.05 mg/l anisomycin (ANS) for 15 minutes. **h,** Global protein synthesis measurements by OPP labeling in control hiPSCi and/or after treatment with 6.25, 12.5, and 25 µM 4EGI-1 for 1 day. Median fluorescence intensity was quantified by flow cytometry (>10,000 cells/ analysis) and normalized to the average value in controls (*n*=2 biological replicates).

## Methods

### Cell culture and hiPSCi differentiation

The reference HPSI0214i-kucg-2 cell line was engineered to express a KRAB-dCas9 from a doxycycline-inducible promoter at the *AAVS1* locus using pAAVS1-PDi-CRISPRnpAAVS1-PDi-CRISPRn (a gift from Bruce Conklin; Addgene plasmid #73500; http://n2t.net/addgene:73500; RRID: Addgene_73500)^26,29^. The HEK293i cell line was constructed in a similar fashion^131^. hiPSCi were cultured in mTeSR Plus on Matrigel-coated plates at 37°C/5% CO_2_. The medium was exchanged every other day, and cells were propagated as clusters with 0.5 mM EDTA/PBS every five days in a ratio of 1:20. For experiments, cells were singularized with Accutase and resuspended in mTeSR Plus containing 10 µM of the ROCK inhibitor Y-27632. Viable cells were counted with Trypan Blue in the Cell Countess II system and pelleted for 5 min at 200 x *g*. Cells were seeded in mTeSR Plus with 10 µM Y-27632, which was removed from the medium on the next day. The hiPSCi line was differentiated into cardiomyocytes, NPC and neurons using small molecules as previously described^29,132–134^. CRISPRi HEK293 cells and Lenti-X™ 293T cells (#632180, Takara) were cultured in DMEM high glucose medium supplemented with 10% FCS at 37°C/5% CO_2_ and passaged using 0.25% Trypsin/EDTA every other day in a ratio of 1:10 to 1:20.

### CRISPRi library design

An adapted version of the CRISPRiaDesign workflow^30^ (https://github.com/mhorlbeck/CRISPRiaDesign) was used to design sgRNAs to target the TSS of selected genes. Modifications were made to the prediction pipeline to incorporate information about SNPs in the HPSI0214i-kucg-2 genome, and to utilize public FAIRE-seq and DNase-seq data for improved sgRNA prediction accounting for chromatin topology^29^. Briefly, variant sites from the HPSI0214i-kucg-2 genome haplotype calls were extracted using gvcftools *extract_variants* v0.17.0, and the resulting genomic variant call format (GVCF) file was indexed using GATK *GenotypeGVCFs* v4.1.0.0. GATK *SelectVariants* v4.1.0.0 with the *-select-type SNP* parameter was used to retain SNP variants only. By creating a sequence dictionary of the reference GRCh37 genome with Picard *CreateSequenceDictionary* v2.17.10, we could replace the identified SNPs by supplying the SNP VCF to GATK *FastaAlternateReferenceMaker* v4.1.0.0. DNase-seq (ENCFF899UPI) and FAIRE-seq (ENCFF000TJL) chromatin accessibility bigWig files downloaded from the ENCODE projects ENCSR794OFW and ENCSR000DCD, respectively, were used in model training to improve accuracy of sgRNA predictions. Using our custom genome, and supplying the H1-hESC chromatin accessibility data and other supplied training data from the CRISPRia pipeline, including sgRNA activity scores and TSS predictions (https://github.com/mhorlbeck/CRISPRiaDesign/tree/master/data_files), an elastic net linear regression model for sgRNA activity predictions was trained. After activity score prediction for each sgRNA, off-targets were predicted per sgRNA as described^30^. The sgRNAs with highest predicted activity scores and no predicted off-targets were used for sgRNA library design.

### Pooled CRISPRi screens

The sgRNA pool for the CRISPRi screens was ordered with matching overhangs for a mU6-sgRNA-EF1-puromycin-mKate2 construct, amplified using KAPA HiFi Hotstart polymerase with the High-Fidelity buffer (95°C/3min - [98°C/20 sec - 56°C/15 sec - 72°C/15 sec] x12 – 72°C/1 min) and cleaned up using the Zymo DNA Clean & Concentrator Kit. The acceptor plasmid was digested with BstXI and BlpI. Gibson assembly reactions were performed in a 3:1 ratio (insert:backbone) and transformed into MegaX competent cells. Colonies were scraped off plates and the plasmid DNA was extracted by Plasmid Plus Midi Kit (Qiagen, #12943). For lentivirus production, Lenti-X 293T cells were co-transfected with the sgRNA plasmid pool and third generation lentiviral packaging plasmid mix (4:1:1, pMDLg/pRRE, Addgene plasmid #12251, pRSV-Rev (Addgene plasmid #12253) and pMD2.G (Addgene plasmid #12259; all a gift from Didier Trono) in a 1:1 ratio with TransIT®-Lenti Transfection Reagent (Mirus, # MIR6603) following the manufacturer’s instructions. After two days, the viral supernatant was collected, filtered through a 0.45 μm PVDF syringe filter and precipitated with Lentivirus precipitation solution (Alstembio, #VC125) at 4 °C overnight. Virus stocks were resuspended in cold PBS and stored at -80 °C.

Growth screens were performed on dividing cells in three biological replicates (hiPSCi, HEK293i and NPC). hiPSCi were transduced by co-seeding cells and viral stocks at the same time in the absence of doxycycline with an initial transduction rate of 30%. Cells were reseeded after two days in medium containing 2 μg/ml puromycin and selected for two passages. After 3 days recovery without puromycin, 3×10^6^ cells representing a 1000x coverage of the sgRNA library, were seeded with and without 2 μM doxycycline and passaged every five days. hiPSCi were harvested after ten days, which corresponds to approximately ten cell doubling times. For HEK293i, 8 μg TransduceIt (Mirus, # MIR6620) was added during transduction, and cells were selected in 1 μg/ml puromycin. HEK293 cells were passaged every three days during the screens and harvested after twelve days, which corresponds to approximately ten cell divisions. 8×10^6^ NPCs were transduced in the presence of doxycycline, reseeded in 2.5 μg/ml puromycin the next day, and selected for two days. The culture was then split in two, and 3×10^6^ cells were seeded either with or without 2 μM doxycycline. NPC were passaged every five days and harvested after 15 days, which corresponds to approximately ten cell doubling times.

Differentiation screens were performed by inducing KRAB-dCas9 expression at the start of the derivation of CM or NPC from hiPSCi, and of neurons from NPC. For survival screens in CM and neurons, KRAB-dCas9 expression was induced at day 15 of CM derivation from hiPSC, and at day 21 of neuron derivation from NPC. Virus transductions were performed in two biological replicates for CM and three biological replicates for neurons. For CM, hiPSCi were transduced and selected with puromycin as for the growth screens, and CM derivation was initialized after hiPSCi selection. CM were split at the reseeding step to two wells either without or with 2 μM doxycycline and cultured for additional 20 days before harvesting. For neurons, NPC were transduced and selected with puromycin as for growth screens, and neuron derivation was initialized after NPC selection. Neurons were split at the reseeding step to two wells either without or with 2 μM doxycycline and cultured for additional 20 days before harvesting.

For library preparation, 5×10^6^ cells were harvested per screen and gDNA was extracted using the Nucleospin Blood Kit (Macherey&Nagel). 20 μg of DNA was amplified using NEBNext Ultra II Q5 Mix. Two different primer sets were used to maximize read variety during library sequencing. Each PCR reaction contained 5 μg DNA and was amplified with the standard program (98°C/2min - [98°C/10sec - 60°C/30sec - 65°C/45sec] x22 - 65°C/5min). PCR reactions for the same sample were pooled and concentrated using the Zymo DNA Clean and Concentrator Kit. The reaction was run on an 8% PAA gel and the PCR product corresponding to the sgRNA library was excised. The gel slices were crushed with a disposable pestle and DNA was eluted in water overnight at room temperature on a rotating wheel. The next day, gel debris was removed by Spin-X filter and the DNA was recovered by ethanol precipitation. The final library concentration was measured with the Qubit dsDNA HS kit. Libraries were sequenced on an NextSeq 500 platform (Illumina).

### Single and dual sgRNA knockdown experiments

Single guide RNA (sgRNA) knockdown experiments were performed in a mU6-sgRNA EF1α-Puro-T2A-GFP construct^29^. For dual sgRNA knockdown, we adapted the construct described in Tzelepis et al.^135^). sgRNAs were ordered with overhangs to the backbone and cloned by Gibson Assembly. Lentiviral transduction was performed as during the pooled screens. For growth assays, the cell population was split into two wells, with and without 2 µM doxycycline, and cultured for a time period of 20 cell divisions. Cells were passaged every four days and the percentage of GFP+ cells was measured on the Attune NxT system. For follow-up analysis, hiPSCi and HEK293i were selected with 2 or 1 μg/ml puromycin for 3-5 days (until the fraction of GFP+ cells exceeded 80%). NPC were selected with 2.5 μg/ml puromycin -/+ Dox, respectively, until >80% cells were GFP+.

### Stalling readthrough reporter experiments

We adapted the previously described reporter constructs^63,73^ by exchanging the promoter CMV promoter with a UbC promoter and replacing the fluorescent cassettes to fit our experimental set-up (GFP to BFP, and RFP to mOrange). We also changed the first 2A skipping sequence to T2A. For this, the UbC-BFP-T2A-FLAG-XBP1stall-P2A-mOrange cassette^63^ was synthesized by Twist Bioscience and inserted into a plasmid containing sequences for lentiviral packaging (Addgene #60955). The XBP1 sequence was removed to yield the “no stall” reporter (Addgene #105686) or replaced with an AAA-encoded poly-lysine sequence (AAA_20_, Addgene #105688). For stalling readthrough assays, puromycin-selected cells transduced with individual sgRNA expression vectors (mU6-sgRNA_EF1A-Puro-GFP) were cultured with 2 µM doxycycline for 3 days, transduced with the stalling constructs, and allowed to recover for 2 days before analysis. Cells were harvested and analysed on the Attune NxT flow cytometer.

### ZNF598^RING^ and ZNF598^WT^ expression

The cDNA for wild-type ZNF598 or the ZNF598^RING^ C29S/C32S mutant^62^ with a C-terminal HA tag was synthesized by Twist Bioscience as an A2UCOE-EF1a-ZNF598-2A-BLC cassette and inserted into a plasmid containing sequences for lentiviral packaging (Addgene #60955). Cells transduced with the expression constructs and cultured for a total of two days before harvesting. For ribosome loading inhibition experiments in hiPSCi, 5 µM 4EGI-1 (Sigma Aldrich #324517) was added for 18 h before harvesting.

### Global protein synthesis measurements

For puromycin-selected cells transduced with mU6-sgRNA_EF1A-Puro-GFP, cells were washed twice with PBS and incubated in methionine-free DMEM supplemented with 10% dialyzed FCS (#A3382001, Thermo Fisher) for 30 minutes. Cells were then labeled with 2 µM L-Homopropargylglycine (HPG, #CLK-1067, Jena Bioscience) for 30 minutes. Control and ZNF598^RING^-expressing hiPSCi were labeled by addition of 20 µM O-propargyl-puromycin (OPP, #C10459, Thermo Fisher Scientific) to culture medium for 30 minutes. Cells were fixed for 10 minutes in 3.7% formaldehyde in TBS, washed with TBS and permeabilized for 15 minutes in TBS-T (0.5% Tween 20). HPG and OPP were labeled with click chemistry by a 30-minutes incubation in 100 mM Tris pH=8, 1 mM CuSO4, 20 µM AF647-Picolyl-Azide, #CLK-1300 (Jena Bioscience), 100 mM ascorbic acid). Cells were washed three times in TBS-T (0.2% Tween) and fluorescence signal was analysed on an Attune NxT flow cytometer.

### Cell cycle analysis by 5-ethynyl-2’-deoxyuridine (EdU) staining

Culture medium was exchanged with fresh medium two hours prior to harvesting. Cells were labeled using the EdU Flow Cytometry kit (Thermo Fisher Scientific, # C10634). In brief, cells were incubated with 10 µM 5-ethynyl-2’-deoxyuridine (EdU) for 1 hour and collected by trypsinization. Cells were washed once in 1% BSA/ PBS and fixed in click-it fixative for 15 minutes. Cells were washed once in 1% BSA/ PBS and permeabilized in the permeabilization and wash reagent for 15 minutes, prior addition of the click it reaction. Cells were incubated for 30 minutes in the dark and washed twice with the permeabilization and wash reagent. Finally, 1 µl of FxCycle dye (Thermo Fisher Scientific, #F10347) was added, cells were incubated for 30 minutes in the dark and fluorescence intensity was measured on an Attune NxT flow cytometer.

### RNA isolation

Culture medium was aspirated and cells were directly lysed in culture plates by adding LiDS/LET buffer (5% LiDS in 20 mM Tris, 100 mM LiCl, 2 mM EDTA, 5 mM DTT, pH7.4, 100 μg/ml Proteinase K). Lysates were collected, incubated at 60°C for 10 minutes, titurated 10 times through a 26G needle, and mixed by vortexing. RNA was extracted with phenol/ 1-Bromo-3-chloropropane (BCP). For this, two volumes of cold acid phenol (pH 4.3), 1/10 volume BCP and 50 µg glycogen (Thermo Fisher Scientific, #AM9510) were added and samples were mixed by vortexing. Phases were separated by centrifugation for five minutes at 10,000 x *g* at 4°C. The aqueous phase was transferred to a new tube and a second phenol/BCP extraction was performed. RNA was precipitated with 100% ethanol (2.5 volumes) at -20°C for 30 min. After 30 minutes of centrifugation at 16,000 x *g* at 4°C, cell pellets were air-dried, and resuspended in RNase-free water. RNA concentration was measured with Nanodrop and samples were stored at -80°C.

### Quantitative RT-PCR

5 μg of total RNA were digested for 30 minutes at 37°C/1500 rpm using 2 U of Turbo DNase (Thermo Fisher Scientific, #AM2238). The RNA was then cleaned up by phenol/ BCP extraction. For the reverse transcription, 1 μg of DNase treated RNA was transcribed to cDNA using the Protoscript II First Strand cDNA Synthesis Kit (NEB, #E6560). Quantitative RT-PCR was performed with the KAPA SYBR Fast qPCR Mix (Roche, #KK4601) and a 1:80 dilution of the cDNA. For analysis, ΔΔCt values were calculated relative to a non-targeting control. qPCR primer efficiency was tested with a serial dilution; only primer pairs with an efficiency between 1.9-2.1 were used for experiments.

### Immunoblotting

Cells were quickly rinsed with cold PBS and lysed by addition of RIPA buffer (20 mM Tris pH7.5, 150 mM NaCl, 1% NP-40, 0.5% sodium deoxycholate, 0.1% SDS) supplemented with 10 µg/ml aprotinin, 20 µM leupeptin, 2.5 µM pepstatin A, 0.5 mM AEBSF and 1x Phosphatase Inhibitor Cocktail (Cell Signaling, #5870) to culture plates. Extracts were collected and incubated on ice for 20 minutes. Insoluble fractions were pelleted by 5-minute centrifugation at 10 000 x *g*/ 4°C. EDTA was added to a final concentration of 5 mM to the supernatants, and protein concentration was then quantified with the Pierce™ BCA Protein Assay Kit (Thermo Fisher Scientific, #23225). 15 μg total protein from each sample were resolved on 4%–12% Bis-Tris precast polyacrylamide gels (Life Technologies) in Bolt™ MOPS SDS Running Buffer (Invitrogen, #B0001). Proteins were then transferred to a 0.2 μM nitrocellulose membrane (Amersham, #10600015) for 30-40 minutes at 25 V using the semi-dry TransBlot Turbo System (Biorad). For total protein visualization, the membranes were stained with Ponceau S solutions (0.5% Ponceau S, 1% acetic acid) for 3 min at room temperature, rinsed with distilled water and imaged on BioRad Imager. Membranes were destained with PBST (0.1% Tween-20), blocked for an hour in 5% milk/PBST (0.1% Tween-20), and further incubated with primary antibodies in blocking solution overnight at 4°C (ZNF598, 1:1000, #ab135921 Abcam; PELO F-4, 1:1000 #sc-393418 Santa Cruz Biotechnology; HBS1L, 1:1000 #HPA029729 Atlas Antibodies; ASCC3, 1:1000 #A304-015A Bethyl Laboratories; eIF2α, 1:1000, #9722 Cell Signaling; eIF2α-p S51, 1:1000, # ab32157 Abcam; p38, 1:1000, #9212 Cell Signaling; p38-p Thr180/Tyr182, 1:1000, #9211 Cell Signaling; eS10 1:1000, #ab151550 Abcam; uS10, 1:1000, #ab133776, Abcam; uS5, 1:1000, #A303-794A Bethyl Laboratories; uS3 (1:1000, #A303-840A Bethyl Laboratories). The next day, membranes were washed in PBST (0.1% Tween-20) and incubated with HRP-labeled secondary antibodies in 5% milk/PBST (0.1% Tween-20) at room temperature for 1 h (anti-rabbit IgG-HRP, 1:4000, #111-035-003 Dianova or anti-mouse IgG-HRP, 1:4000, #115-035-003 Dianova). Membranes were incubated for five minutes in SuperSignal West Pico PLUS (Thermo Fisher Scientific, #34577) and proteins were visualized on an iBright system (Thermo Fisher Scientific).

For comparisons of basal stress activation in hiPSCi and HEK293i, membranes stained with eIF2α-p and p38-p antibodies were stripped using Restore Western Blot stripping buffer (Thermo Fisher Scientific, #21059) for 30 minutes at room temperature, blocked for another hour, and re-probed with eIF2α and p38 antibodies, respectively. For polysome experiments, membranes stained with eS10 and uS10 antibodies were re-probed with uS5 and uS3 antibodies, respectively, which had different molecular weights to distinguish the potential remaining antibody signal from the primary staining.

### Immunostaining

Cells were seeded on glass bottom dishes (ibidi, #80827) and fixed in 3.7% formaldehyde for 10 minutes at room temperature. Formaldehyde was stepwise replaced with PBST (0.02% Tween-20). hiPSCi and NPC were then permeabilized for 10 minutes with 0.5% Triton-X100 in PBST (0.02% Tween-20), and blocked for one hour in blocking solution (3% BSA/ 0.1% Triton-X100 in PBS). Cells were incubated overnight with the primary antibody diluted in blocking solution at 4°C (POU5F1 C-10 (1:400, SCBT, #sc-5279), NANOG P1-2D8 (1:200, Millipore, #MABD24), PAX6 (1:200, Abcam, #ab5790), NES (1:200, R&D Systems, #MAB1259). Cells were washed three times in PBST (0.02% Tween-20), and incubated with 500 ng DAPI (DAPI, #10236276001, Roche) and the secondary antibody diluted in blocking solution for one hour at room temperature (goat anti-mouse Alexa Fluor 488, #A-11001, Thermo Fisher Scientific, 1:2000; goat anti-rabbit Alexa Fluor 488, #A-11034, Thermo Fisher Scientific, 1:2000; goat anti-mouse Alexa Fluor 633, #A-21052, Thermo Fisher Scientific, 1:500). Cells were washed three times in PBST (0.02% Tween-20) before imaging. Neurons were permeabilized for 10 minutes in PBST (0.7% Tween-20), and blocked for one hour in blocking solution (1% BSA/ 0.1% Triton-X100/ 10%FCS in PBS). Cells were washed once in 0.1% BSA/PBS and incubated overnight with the primary antibody diluted in 1% BSA/ PBS at 4°C (MAP2, #ab92434, Abcam, 1:1000; CHAT, #ab6168, Abcam, 1:200). Cells were washed three times in 0.1% BSA/ PBST (0.05% Tween) and incubated with 500 ng DAPI and the secondary antibody diluted in 1% BSA/ PBS for one hour at room temperature (goat anti-rabbit A633, #A-21070, Thermo Fisher Scientific, 1:500; goat anti-chicken A488, #A-11039, Thermo Fisher Scientific, 1:2000). Cells were washed three times in 0.1% BSA/ PBST (0.05% Tween) before imaging. Cardiomyocytes were blocked and permeabilized in 3 % BSA, 0.1 % Triton-X in PBS for one hour at room temperature. Cells were washed three times with PBST (0.1 % Tween), and incubated with primary antibody (anti-ACTN2, Sigma Aldrich #A7811, 1:800; anti-cTNT, CT3, deposited to the DSHB by Lin, J.J-C., 1:5) diluted in blocking solution at 4°C overnight. Cells were washed again three times with PBST (0.1 % Tween) and incubated with 500 ng DAPI and the secondary antibody (anti-mouse Alexa Fluor 488, 1:2000) diluted in blocking solution for one hour at room temperature in the dark. Cells were washed three times in PBST (0.1 % Tween) before imaging.

### Flow cytometry analysis of eIF2α and p38 phosphorylation

Culture medium was exchanged with fresh medium two hours prior to harvesting. Cells were collected by trypsinization, washed once with 1% BSA/ PBS, and fixed by slowly adding 3.7 % formaldehyde in PBS and incubated for 10 minutes at room temperature. Cells were washed once with 1% BSA/ PBS and incubated with ice-cold methanol for 10 minutes. Cells were washed once with 1% BSA/ PBS and incubated with the primary antibody (eIF2α-p S51, 1:100, # ab32157 Abcam; p38-p Thr180/Tyr182, 1:100, #9211 Cell Signaling) for 45 minutes at room temperature. Cells were washed twice with 1% BSA/ PBS and incubated with the secondary antibody (anti-rabbit Alexa Fluor 633, 1:200, Thermo Fisher Scientific, #A21070) for 45 minutes at room temperature. Cells were washed three times with 1% BSA/ PBS, resuspended in PBS and fluorescence intensity was measured on an Attune NxT flow cytometer.

### Inhibitor treatments

To pharmacologically block the ISR and RSR, cells were treated with chemical inhibitors at concentrations >-fold higher than their respective IC50 in human cells^82,136^. 1.25 µM GCN2i (A-92, Axon Medchem, #Axon1386), 4 nM PERKi (GSK2606414, Merck, #516535), 1 µM p38i (SB203580, Cell Signaling, #5633), or 50 nM ISRIB (Sigma Aldrich, #SML0843) were added simultaneously with doxycycline to sgRNA-transduced cells. Fluorescence intensity was measured on an Attune NxT flow cytometer. To induce p38 phosphorylation, cells were treated with 0.05 mg/l anisomycin (ANS, Sigma Aldrich, #A9789) for 15 minutes prior to harvesting. To induce ER stress and eIF2α phosphorylation, cells were treated with 2.5 µM tunicamycin (TM, Sigma Aldrich, #T7765) for 2 hours prior to harvesting.

### Polysome profiling and protein analysis of polysome fractions

Cells were grown in a 10-cm dish until ∼80 % confluency and cell medium was exchanged two hours prior to harvesting. For ribosome run-off experiments, cells were treated for 2.5 min with 2 µg/ml homoharringtonine (HAR, Cayman Chemical, #15361) prior to harvesting. To induce ribosome collisions, cells were treated with 0.05 mg/l anisomycin (ANS, Sigma Aldrich, #A9789) for 15 minutes prior to harvesting. Cells were quickly washed with ice-cold PBS supplemented with 10 mM MgCl_2_ and 100 µg/ml CHX, and snap-frozen in liquid nitrogen. Cells were harvested in lysis buffer containing 50 mM HEPES pH 7.4, 150 mM KCl, 15 mM MgCl_2_, 1% Triton-X 100, 1mM DTT, 1x protease and phosphatase inhibitors (10 µg/ml aprotinin, 20 µM leupeptin, 2.5 µM pepstatin A, 0.5 mM AEBSF and 1x Phosphatase Inhibitor Cocktail (Cell Signaling, #5870)), 1 mM TCEP, and 10 mM NEM. Cells were directly loaded on 10-50% sucrose gradients in 50mM HEPES pH 7.5, 150mM KCl, 15 mM MgCl_2_, 1mM DTT, 1 mM TCEP, and 1.2 ml fractions were collected. Protein was extracted by TCA/ acetone and ethanol precipitation. 5 % sodium deoxycholate was added to each fraction and incubated on ice for 30 minutes. 20 % TCA was added in a 1:1 ratio vortexed and incubated for another 30 minutes on ice. Protein pellets were collected by centrifugation for 30 minutes at 13 000 x *g* at 4°C. Pellet was washed once with 100% acetone and dissolved in 4x Laemmli sample buffer. One-quarter of each fraction was loaded per well.

### RNA-Seq library construction

250 ng of total RNA were used for library preparation with the Universal Plus RNA-Seq with NuQuant kit (Tecan, #0361). The fragment size of final libraries was determined on an Agilent TapeStation and the concentration was quantified using the Qubit dsDNA HS assay. 80-bp single-end sequencing was performed on a NextSeq 500 platform (Illumina).

### Ribosome profiling library construction

Ribosome footprint libraries were prepared as previously described^55,137^ with minor modifications. Cells were grown in a 10-cm dish until ∼80 % confluency and cell medium was exchanged two hours prior to harvesting. Cells were quickly washed with ice-cold PBS supplemented with 10 mM MgCl2 and 100 µg/ml CHX, and snap-frozen in liquid nitrogen. Cells were thawed on ice and scraped off the plate in 400 µl polysome lysis buffer (20 mM Tris pH=7.4, 150 mM NaCl, 5 mM MgCl_2_, 1% Triton-X100, 1 mM DTT, 100 µg/ml CHX, 25 U/ml Turbo DNase (Thermo Fisher Scientific, #AM2238), 0.1% NP-40. Samples were vortexed vigorously, triturated through a 26G gauge needle, and spun down for 7 minutes at 16 000 x *g*/ 4°C. Supernatant was transferred to a new tube and RNA concentration was measured with the Qubit RNA HS Kit. Aliquots of 20 µg RNA in 200 µl polysome lysis buffer were snap-frozen and stored at -80°C. 20 µg RNA in 200 µl polysome lysis buffer were digested with 50 U RNase I (Thermo Fisher Scientific, #AM2295) for 45 minutes at 2 000 rpm/22°C.

RNase I digestion was stopped with 100 U Superase In (Thermo Fisher Scientific, #AM2694), mixed by pipetting and extracts were loaded on a 1 M sucrose cushion. For this, 0.9 ml sucrose in polysome lysis buffer was carefully inserted below the 200-µl digested extract, followed by centrifugation for 75 minutes at 120,000 rpm/4°C in a S120AT2 rotor (Thermo Fisher Scientific). The pellet was dissolved in 400 µl LiDS/LET lysis buffer and RNA was isolated by phenol extraction (see *Total RNA isolation*). 3 µg of RNA were mixed with 2x RNA loading dye (45% formamide, 0.25x TBE, 0.025 % SDS), boiled for three minutes at 90°C and loaded on 15% polyacrylamide/7M urea/1×TBE gels. Fragments in the range of 19 to 32 nucleotides (monosome footprint libraries) and 50 to 80 nucleotides (long footprint libraries) were excised from the gel based on size markers and gel slices were crushed with a disposable pestle. RNA was eluted from gel slices with 400 µl gel elution buffer (0.3 M NaOAc pH 4.5, 0.25% SDS, 1 mM EDTA pH=8.0) by incubation for 10 minutes at 65°C, followed by snap-freezing on dry ice for 10 minutes, thawing for 5 min at 65°C, and an overnight elution at room temperature on a rotating wheel. Gel debris was removed by a Spin-X filter (Corning) and RNA was purified by ethanol precipitation. Size-selected RNA was dephosphorylated for 45 minutes at 37°C using 5 U of T4 PNK (NEB, #M0201S) and directly mixed with pre-adenylated adapters containing 5 random nucleotides at their 5’ ends^137^ in 1x T4 RNA ligase buffer (25% PEG-8000, 20 U Superase In and 200 U T4 RNA Ligase 2, truncated KQ (NEB, #M0373S)). The linker ligation mix was incubated for 3 hours at 25°C, and products were pooled and concentrated with the Oligo Clean and Concentrator kit (Zymo, #D4060). Ligation products were then size selected on a 12% polyacrylamide/7M urea/1×TBE gel and purified as above in gel elution buffer. The sample concentration was quantified with the Nanodrop and 50 ng of the linker-ligated sample was used for rRNA depletion using the Ribo-Seq riboPOOL h/m/r depletion kit (siTOOLs, dp-K024-000050). The footprints were annealed with RT primer (5’-pRNAGATCGGAAGAGCGTCGTGTAGGGAAAGAG/iSp18/GTGACTGGAGTTCAGACGTGTGCTC-3’) at 65°C for 5 minutes and subsequently reverse transcribed for 30 minutes at 50°C in 1x Protoscript II Buffer containing 0.5 mM dNTPs, 10 mM DTT, 20 U Superase In and 200 U Protoscript II (NEB, #E6560S). After RT, the RNA was hydrolysed by addition of 0.1 M NaOH and incubation for 5 minutes at 90°C. cDNA products were size selected on a 12% polyacrylamide/7M urea/1×TBE gel. The DNA was eluted from crushed gel slices for 60 minutes at 1500 rpm/70°C in 1x TE buffer. The crushed gel was removed by filtering samples through a Spin-X filter (Corning) and RNA was purified by ethanol precipitation. The gel-purified RT product was circularized by addition of 3 µM recombinant TS2126 RNA ligase 1 (commercially sold as CircLigase)^138^ in circularization buffer (50 µM ATP, 2.5 mM MnCl_2_, 50 mM MOPS, pH=7.5, 10 mM KCl, 5 mM MgCl_2_, 1 mM DTT and 1 mM Betaine) and incubation for three hours at 60°C, followed by heat inactivation for 10 minutes at 80°C. Libraries were amplified from circularized cDNA using a common forward primer (5’-AATGATACGGCGACCACCGAGATCTACACTCTTTCCCTACACGACGCT*C-3’), an index-containing reverse primer (5’-CAAGCAGAAGACGGCATACGAGAT<u>NNNNNN</u>GTGACTGGAGTTCAGACGTGT*G-3’), and the KAPA HiFi DNA Polymerase (Roche) in 1X HiFi buffer. Samples were initially denatured at 95°C for 3 min, followed by twelve cycles of 98°C for 20 s, 62°C for 30 s, 72°C for 15 s at a ramp rate of 3°C/sec. PCR products were size selected on an 8% polyacrylamide/1 × TBE gel. The DNA was eluted from crushed gel slices on a rotating wheel overnight at room temperature in DNA elution buffer (300 nM NaCl, 10 mM Tris-Cl pH=7.5, 0.2% Triton-X 100). Gel debris was removed by filtering samples through a Spin-X filter (Corning) and RNA was purified by ethanol precipitation. Libraries were quantified with the Qubit dsDNA High Sensitivity kit and 75 - 86-bp single-end sequencing was performed on a NextSeq 500 platform (Illumina).

### CRISPRi screen data analysis

sgRNA-Seq datasets were processed following the ScreenProcessing pipeline (https://github.com/mhorlbeck/ScreenProcessing)^30^. Gene-avegared sgRNA log_2_ fold changes were calculated by normalizing sgRNA log_2_ enrichment of the average top three sgRNAs using *calculate_ave = True* and *best_n = 3*. The Mann-Whitney *p*-value was calculated using the average log_2_ fold change from all nine sgRNAs targeting the same gene TSS compared to non-targeting controls by setting *calculate_mw = True*. For defining gene essentiality, gene-averaged log_2_ fold changes of genes with two alternative TSS were collapsed by choosing the stronger magnitude effect with a Mann-Whitney *p*-value ≤ 0.1. For comparison of shared essential genes, datasets from CRISPR-inferred common essential genes in DepMap 23Q4 and genome-wide CRISPRi screens in WTC11 hiPSC^34^ and in WTC11 hiPSC-derived neurons^35^ (FDR ≤ 0.1) were subset for the 262 genes targeted in our screens.

### RNA-Seq data analysis

Potential 3’ adapters in RNA-Seq reads were first automatically detected and removed using TrimGalore v0.6.4. Using default settings, reads with length ≥20 nt were retained. Alignment of trimmed reads to the GRCh38 genome was performed with STAR v2.6.1c to an index built including the GENCODE comprehensive annotation gtf file in order to guide splice-aware alignment. The following parameters were used for alignment with STAR to ensure no more than one mismatch per read and retention of only uniquely mapped reads: *--outSAMtype BAM SortedByCoordinate --outFilterMultimapNmax 1 --outFilterMismatchNmax 1.* Read counts per gene were then generated using featureCounts v1.6.2 and only the protein-coding gene annotation subset from GENCODE. This count table was then used as input for DESeq2 v1.38.1 to perform differential gene expression analysis. DESeq2 was run with default settings to quantify differential expression between conditions. Using the resulting log_2_ fold-changes, heatmaps comparing gene differential expression between conditions were assembled using ComplexHeatmap v2.14.0^139^, where only significant values were retained (*p-adj* ≤ 0.05). Gene ontology enrichment was analyzed of significantly altered log_2_ fold-changes (*p-adj* ≤ 0.05) using ClusterProfiler v4.4.4^140^ with a significance cut-off of ≤ 0.01 (Benjamini-Hochberg method).

### Ribosome profiling data analysis

A custom reference transcriptome annotation was built for the GRCh38.p13 genome based on the Matched Annotation from the <u>NCBI</u> and <u>EMBL-EBI</u> (MANE) and the best scoring transcript annotated in APPRIS (https://apprisws.bioinfo.cnio.es/pub/current_release/datafiles/homo_sapiens/GRCh38/appris_data.appris.txt).^141^ Sequencing libraries were demultiplexed and adapter-trimmed with Cutadapt v2.5. Indels in the alignment to the adapter sequence were disabled with *--no-indels* and low-quality bases were removed from both 5’ and 3’ ends with *-q 30,30*. Reads without adapters were discarded with *--trimmed-only*. Reads were further trimmed to remove the two 5′-RN nucleotides introduced by circularization from the RT primer with *-u 2*. Trimmed reads longer than 10 nt were aligned to a human ribosomal RNA reference using Bowtie v1.2.2^142^ with the following options: *-p 40 -S --best*. rRNA-filtered reads were aligned to GRCh38.p13 using STAR v2.6.1c^143^ with the following options: *--outFilterMultimapNmax 1 --outSAMtype BAM SortedByCoordinate --outFilterMismatchNmax 0 --alignEndsType Local -- seedSearchStartLmax 14 --alignIntronMax 10000 --outFilterIntronMotifs RemoveNoncanonicalUnannotated --quantMode TranscriptomeSAM --outSAMattributes NH HI AS nM NM MD.*The A-site codon in each mapped read was identified with Scikit-ribo, which uses a random forest model with recursive feature selection trained on reads mapping to the start codon^144^. Kallisto 0.44.0 with parameters *-b 100 --single -l 180 -s 20 -t 40* was used to quantify transcript abundances in Transcripts Per Million (TPM) from RNA-Seq data based on MANE and APPRIS annotation.

